# Genomic and phenotypic divergence unveil microgeographic adaptation in the Amazonian hyperdominant tree *Eperua falcata* Aubl. (Fabaceae)

**DOI:** 10.1101/312843

**Authors:** Louise Brousseau, Paul V. A. Fine, Erwin Dreyer, Giovanni G. Vendramin, Ivan Scotti

## Abstract

Plant populations can undergo very localized adaptation, allowing widely distributed populations to adapt to divergent habitats in spite of recurrent gene flow. Neotropical trees - whose large and undisturbed populations often span a variety of environmental conditions and local habitats - are particularly good models to study this process. Here, we carried out a genome scan for selection through whole-genome sequencing of pools of populations, sampled according to a replicated sampling design, to evaluate microgeographic adaptation in the hyperdominant Amazonian tree *Eperua falcata* Aubl. (Fabaceae). A high-coverage genomic resource of ∼250 Mb was assembled *de novo* and annotated, leading to 32,789 predicted genes. 97,062 bi-allelic SNPs were detected over 25,803 contigs, and a custom Bayesian model was implemented to uncover candidate genomic targets of divergent selection. A set of 290 divergence outlier SNPs was detected at the regional scale (between study sites), while 185 SNPs located in the vicinity of 106 protein-coding genes were detected as replicated outliers between microhabitats within regions. Outlier genomic regions are involved in a variety of physiological processes, including plant responses to stress (e.g., oxidative stress, hypoxia and metal toxicity) and biotic interactions. Together with evidence suggesting microgeographic divergence in functional traits, the discovery of genomic targets of microgeographic adaptation in the Neotropics is consistent with the hypothesis that local adaptation is a key driver of ecological diversification, operating across multiple spatial scales, from large- (i.e. regional) to microgeographic- (i.e. landscape) scales.

## Introduction

Plant populations can undergo extremely localized adaptation, allowing populations of widely distributed species to adapt to divergent habitats in spite of recurrent gene flow. This phenomenon has been repeatedly observed in annual plants growing in strongly contrasting yet geographically adjacent habitats, particularly in cases with large differences in soil composition (Antonovics, 2006; Antonovics & Bradshaw, 1970; Bradshaw, 1960; Gould, McCouch & Geber, 2014; Jain & Bradshaw, 1966); in some cases, molecular evidence also supports adaptation to habitat mosaics (Fustier et al., 2017; Turner, Bourne, Von Wettberg, Hu, & Nuzhdin, 2010). In a study of *Howea* palm trees, Savolainen et al. (2006) provided strong evidence of a link between adaptation to habitat mosaics and sympatric speciation. The process underlying these instances of adaptation with gene flow has been dubbed ‘microgeographic adaptation’, whereby adaptive divergence occurs at geographical scales of the same order of magnitude as gene dispersal distance (Richardson, Urban, Bolnick, & Skelly, 2014). Adaptation to multiple optima within a continuous population has been identified as a major mechanism for the maintenance of genetic diversity (Delph & Kelly, 2014). Moreover, conditions under which adaptive divergence can occur with gene flow have been explored theoretically with models including two environmental patches connected by varying levels of gene flow (Bulmer, 1972; Hendry, Day, & Taylor, 2001; Schmid & Guillaume, 2017; Yeaman & Guillaume, 2009; Yeaman & Otto, 2011). These simulations show that a wide range of migration-selection equilibrium states can lead to adaptation to each patch with maintenance of polymorphism (i.e. maintenance of patch specialists, as opposed to the emergence of a generalist genotype).

Data from perennial plants, and from trees in particular, suggest that this may be a general phenomenon, that can be readily observed at the phenotypic level (Brousseau, Bonal, Cigna, & Scotti, 2013; Lind et al., 2017; Vizcaíno-Palomar, Revuelta-Eugercios, Zavala, Alía, & González-Martínez, 2014; Yeaman & Jarvis, 2006) (but see Latreille & Pichot (2017) for a case of a complete lack of microgeographical adaptation). At the molecular level, association between alleles (or genotypes) and habitats has been detected in a variety of studies (see review by Savolainen et al., 2007), suggesting an important role for intra-specific and intra-populational variation in the evolution of habitat specialization (Scotti, González-Martínez, Budde, & Lalagüe, 2016). The advent of genome-wide approaches has increased detection power, allowing to more precisely evaluate the genetic bases of microgeographic adaptation (Eckert et al., 2015; Fustier et al., 2017; Izuno et al., 2017; Lind et al., 2017; Turner et al., 2010). In general, genome scan studies have found only a minority of loci (on the order of a few percent of the total) that exhibit microgeographic disruptive selection. Yet genome-wide, sequencing-based approaches can go beyond a finer estimation of the number of loci involved in the process, because they permit formulating functional interpretations of the divergence process as sequence annotation can suggest putative function of the target loci (Eckert et al., 2015, 2010; Fustier et al., 2017; Turner et al., 2010).

Microgeographic adaptation involves the filtering of phenotypes and genotypes that favor adaptation to habitat patches, and therefore can be viewed as an initial step towards the diversification of populations, ecotypes and eventually species divergence. In particular, environmental filtering has frequently been invoked as a driver of niche partitioning and species diversification in the Neotropics (Baraloto, Morneau, Bonal, Blanc, & Ferry, 2007; Fine, 2015; Fine, Daly, & Cameron, 2005; Fine, Mesones, & Coley, 2004; Fine et al., 2013; Kraft, Valencia, & Ackerly, 2008); intra-populational diversification processes across environmental gradients and/or habitat patches thus may play a key role in the generation of functional diversity. Moreover, species’ short-term adaptive potential (De Kort & Honnay, 2017; Harrisson, Pavlova, Telonis-Scott & Sunnucks, 2014) largely depends on standing genetic variation, and microgeographic adaptation can therefore play a role in the maintenance of adaptive genetic diversity. Indeed, highly heterogeneous landscapes create the conditions for microgeographic adaptations to various environmental conditions and would increase inter-habitat diversity and global diversity in meta-populations considered at higher spatial scales, analogous to beta- and gamma-diversity across species pools in ecology. Measuring the extent of microgeographic adaptation in the wild is thus of major importance, as it could provide insights into the ability of populations to evolve towards new phenotypic and genetic optima under global climate change.

Trees are particularly good models to study microgeographic adaptation: many wild tree species have large populations with widespread distribution spanning a variety of microhabitats, harbor large amounts of genetic diversity, and can disperse over long distances (Fetter, Gugger, & Keller, 2017; Hamrick & Godt, 1990; Neale & Kremer, 2011; Petit & Hampe, 2006; Sork et al., 2013). Trees should therefore not display major departures from the conditions approximating selection-migration equilibrium; nevertheless, their long generation span, combined with continuously changing environmental conditions, may prevent them from actually reaching equilibrium. Moreover, methods to detect locus-specific divergence (and infer selection) should work quite well with microgeographic processes in populations of the most common forest trees, because their relatively shallow (spatial) population structures and large effective population sizes should minimize the introduction of biases (e.g., because of drift) and the generation of false positives.

Lowland Amazonian rainforests, whose landscapes show fine patchiness of contrasting topographic and edaphic conditions varying over microgeographic scales (**Fig. 1a,c,d**) represent an ideal study system to investigate microgeographic adaptation processes. In Amazonian rainforests, the recent study by ter Steege et al. (2013) reported 227 hyperdominant tree species accounting for half of all stems. Some of these widespread species are also ecological generalists, whose preferences could be explained by individual phenotypic plasticity or by microgeographic adaptation to habitat patchiness (Fortunel et al., 2016). We focus here on dense stands of the hyperdominant tree *Eperua falcata* Aubl. (Fabaceae) in French Guiana that occur across environmentally heterogeneous areas, with the goal of identifying genomic loci undergoing divergence at the microgeographic scale between habitats. To do this, a genome scan was carried out through a replicated sampling design combining regional and microgeographic spatial scales (i.e. replicates of microhabitats, ∼300 m apart, in two study sites, ∼300 km apart). We analyzed patterns of genomic and phenotypic divergence from large (i.e. regional) to microgeographic spatial scales. A custom Bayesian framework (Brousseau et al., 2016) was designed to identify locus-specific departures from hierarchical genome-wide divergence in replicated sampling designs. This method was preferred over other outlier detection methods because it is suitable for analyzing Pool-Seq data and it is specifically designed to identify adaptive divergence in nested sampling designs under an explicit hierarchical model of neutral divergence, taking advantage of the cumulative divergence effects occurring in replicated population pairs. Results obtained through the custom Bayesian framework were compared to standard methods for verification. The results of tests for genomic divergence at nested geographical scales were supplemented with trait divergence data, obtained from a large-scale reciprocal transplant experiment involving the very same populations and sites studied at the genomic level. The combination of these two approaches allowed us to formulate detailed interpretations of the observed diversity patterns.

**Fig. 1.**
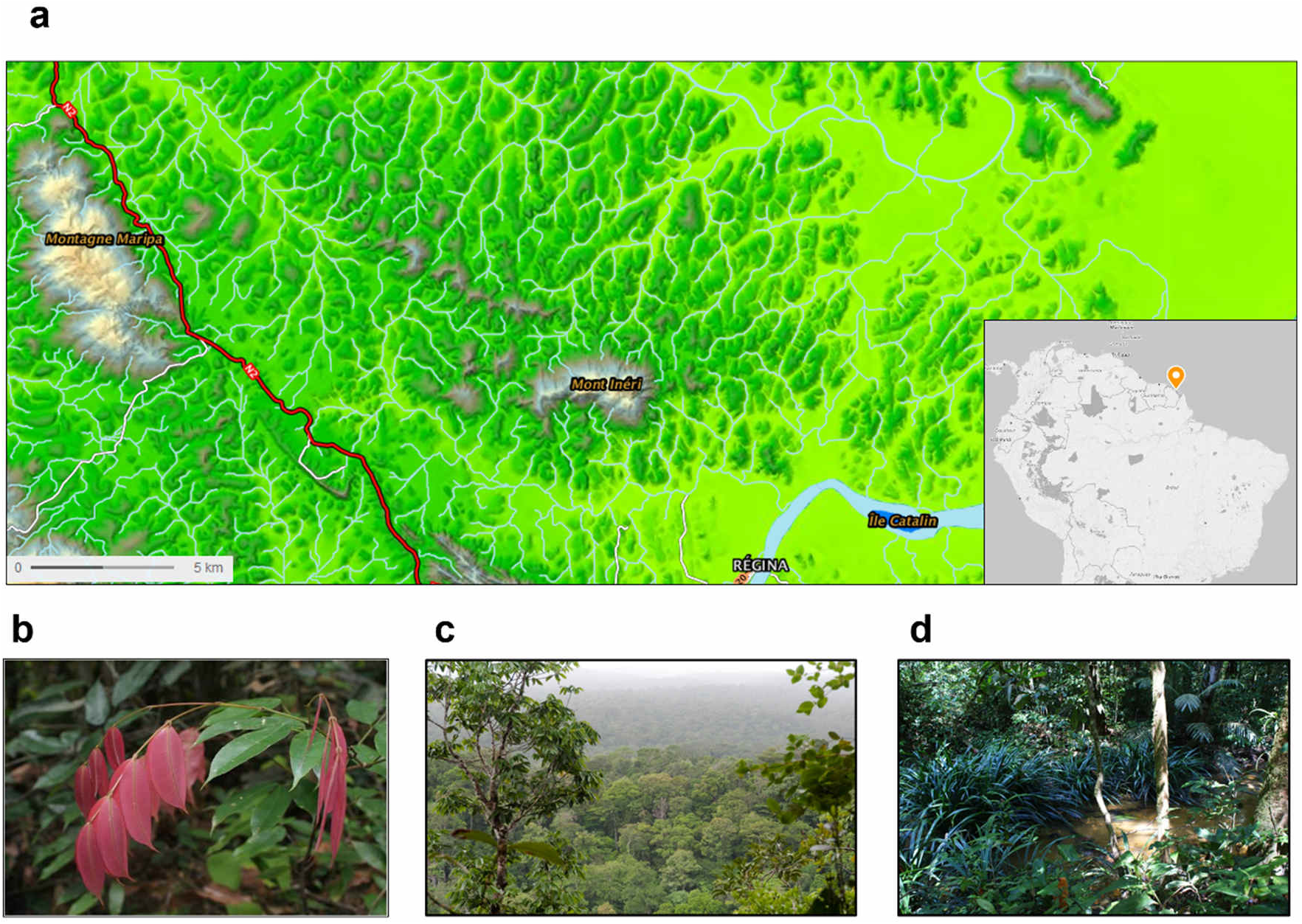
(a) Relief map of forest landscapes in the region of Régina, French Guiana. The map was created through the French ‘Géoportail’ (https://www.geoportail.gouv.fr/): official geospatial data were provided by the French Institute of Geographic and Forest Information ©IGN 2017. (b) *Eperua falcata* sapling. (c) and (d) Forest communities associated with topography and related soil types: *terra-firme* forests on ferralitic soils at the top of hills (c), water-logged forests on hydromorphic soils in bottomlands (d).

## Materials and Methods

### Species model

*Eperua falcata* Aubl. (Fabaceae, Caesalpinioideae, **Fig. S1**) is a reportedly non-nodulating (Villadas, Fernández-López, Ramírez-Saad, & Toro, 2007) tree species. It is widespread in Northeast Amazonia, from North Brazil to Venezuela through the Guiana shield (https://demo.gbif.org/species/2943522), and it is a ‘hyperdominant’ species, the 13^th^ most abundant tree species of the Amazonian tree flora (ter Steege et al., 2013). It is an evergreen, late-successional canopy tree species and it is often emergent. It is shade hemi-tolerant (Bonal, Barigah, Granier, & Guel, 2000) and can tolerate both drought (Bonal, Atger, et al., 2000; Bonal, Guehl, & Champenoux, 2001) and water-logging (Baraloto et al., 2007). Its distribution is very clumped and populations are commonly found at high densities. It is a generalist with respect to microgeographic edaphic conditions, although it is most abundant in water-logged forests of bottomlands (Baraloto et al., 2007). *E. falcata* flowers and fruits during the rainy season, and seeds reach maturity in April-May, synchronously among trees in all microhabitats. Its large pollen grains (>100μm) are dispersed by animals such as bats (Cowan, 1975), while seed dispersal is autochorous: heavy seeds are dispersed around crowns of mother trees through explosive pod dehiscence.

For this species, previous studies have already reported high heritability in quantitative growth and leaf traits in a common garden experiment (Brousseau et al., 2013), suggesting high levels of standing genetic variation for functional phenotypic traits. In addition, seedlings native to different microhabitats displayed genetically-driven differences in phenotypic traits probably due to a combination of neutral processes (i.e. short-distance seed flow and local inbreeding) and local adaptation (Audigeos, Brousseau, Traissac, Scotti-Saintagne, & Scotti, 2013; Brousseau, Foll, Scotti-Saintagne, & Scotti, 2015).

### Sampling design and study sites

We sampled populations that replicated the microgeographic environmental contrast ‘bottomland *versus* terra-firme’ (∼300 m apart) at two regional locations (i.e. study sites) in French Guiana (∼300 km apart), **Fig. 2a**. Laussat (western French Guiana) and Regina (eastern French Guiana) study sites are located on the coastal Guiana Shield and experience different rainfall regimes (∼2,500 and 3,500 mm/year, respectively) (Brousseau et al., 2015). Each regional location harbors different microhabitats (hygromorphic sandy bottomlands and ferralitic *terra-firme* plateaus or hills). In Laussat, a plateau of low elevation is adjacent to a permanently water-logged bottomland, while Regina is composed of higher elevation hilltops bordered by abrupt slopes surrounding a seasonally flooded bottomland forest, **Fig. S2**. *E. falcata* trees are found through the study area, ensuring gene flow between tree populations occupying different microhabitats (Brousseau et al., 2015), **Fig. 2b**. *E. falcata* stem (> 20 cm diameter at breast height) density varies between 30 and 50 trees / ha at these locations (Brousseau et al., 2015). Twenty adult trees for each of the four *regional location × microhabitat* combination were sampled, totaling 80 individuals.

**Fig. 2.**
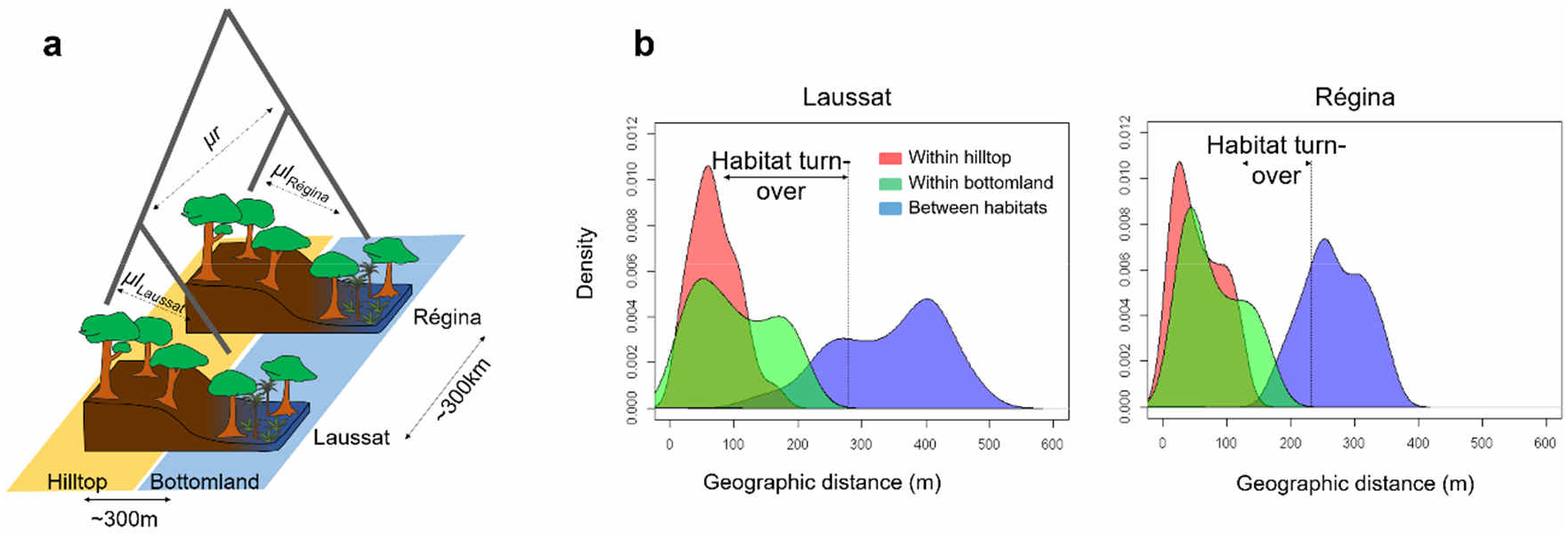
(a) Sampling design scheme and correspondence with genome-wide parameters of divergence estimated through the hierarchical Bayesian model at regional (*μr*) and local scales (*μl*_*Laussat*_ and *μl*_*Régina*_ respectively). (b) Distribution of geographic distances between trees in the study sites of Laussat (Western French Guiana) and Régina (Eastern French Guiana). Vertical dotted bars indicate the range of habitat turnover in each study site.

Details on soil characteristics of the two sites and on growth and functional characters of the sampled trees are reported in **File S1**. Soil properties (including soil type composition, clay/silt/sand composition, fertility, and water table depth) vary between sites and microhabitats, providing a basis for population divergence. The sampled trees have similar diameter at breast height (d.b.h.) and have similar canopy status (almost all are dominant trees) across sites and microhabitats, indicating that sampled populations are demographically homogeneous, although individuals exhibited phenotypic divergence in some traits.

### Molecular methods

Genomic DNA was extracted from fresh leaves according to a modified CTAB protocol (Doyle & Doyle, 1987): CTAB buffer was supplemented in proteinase K in concentrations of 80 μg/ml, and the deproteinization step was realized twice (once with Dichloromethane:Chloroform:Isoamylalcool 24:24:1 and once with Chloroform:Isoamylalcool 24:1). Genomic DNA was eluted in Tris-EDTA 0.5X with RNAse A in concentration 10 μg/ml, and quantified twice using a Nanodrop 2000. Individuals belonging to the same regional location and microhabitat were pooled together in equimolarity, resulting in four libraries of pooled individuals. Sequencing libraries were prepared with a TruSeq Kit (Illumina, San Diego, CA) following manufacturer’s instructions. The four libraries were sequenced (paired-end) on three lanes of an Illumina HiSeq2500. Sequencing library preparation and sequencing were carried out by IGA (Udine, Italy).

### Bioinformatics pipeline and computation of basic diversity statistics

The bioinformatics pipeline is described in **File S2**. Nei’s molecular diversity π (Nei, 1987) was computed for each of the four sub-populations at the contig level with unphased SNP data, under the hypothesis of random mating, as the sum of unbiased estimates of heterozygosity at the nucleotide (SNP) level (Nei, 1987):

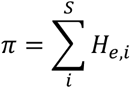

with S = number of polymorphic loci in a contig, and

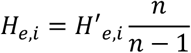

where n = haploid sample size and *H*’_e,i_ = expected heterozygosity at the i^th^ polymorphism in the contig. Because different contigs have different lengths, to compare across contigs we present *π*_*L*_ = *π*/*L*, where L is contig length, that is, molecular diversity per base.

### Bayesian modelling of genomic divergence and outlier detection

A Bayesian modelling approach was used to identify outlier SNPs under selection (Brousseau et al., 2016) has described this Bayesian framework in detail and its power has been assessed through simulations, revealing low false discovery and false non-discovery rates. Briefly, allele frequencies were inferred from read counts and pairwise locus-specific *G*_*ST*_ was estimated in the model, which integrates this way coverage variability across the genome and libraries.

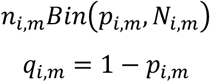

where *n*_*i,m*_ is the read count of the reference allele and *N*_*i,m*_ the total number of read counts at marker *m* in population *i. p*_*i,m*_ and *q*_*i,m*_ are allele frequencies of the reference and the alternative alleles respectively.

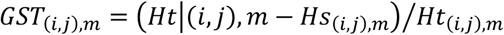

*G*_*ST(i,j),m*_ is the Nei’s fixation index (i.e. genetic differentiation) between populations *i* and *j* at marker *m. H*_*T(i,j),m*_ and *H*_*S(i,j),m*_ are the expected heterozygosities in the total metapopulation and within subpopulations respectively:

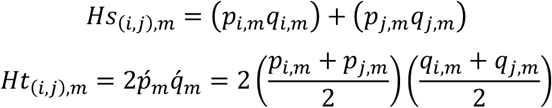

Pairwise *G*_*ST*_ was further partitioned into genome-wide and locus-specific components according to a hierarchical framework: neutral (genome-wide) divergence both between study sites and between microhabitats within each study sites (**Fig. 2a**), and adaptive (locus-specific) divergence both between sites and between microhabitats common to both sites.

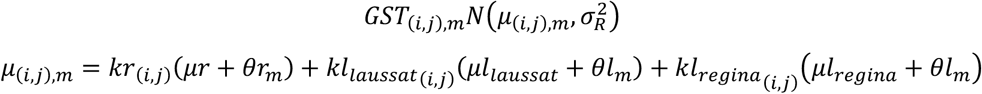

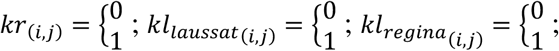

*μr* and *μl* are the global mean of differentiation (over all markers) at regional and local scales respectively: *μr* is the global mean of differentiation between sites, *μl_laussat* is the global mean of differentiation between microhabitats within the study site of Laussat, and *μl_regina* is the global mean of differentiation between microhabitats within the study site of Régina.

*kr*_*i,j*_ and *kl*_*i,j*_ are binary matrices describing whether the populations *i* and *j* belong to the same study site (regional scale) and to the same microhabitat within each study site (local scale): *kr*_*i,j*_ = 1 if the two populations *i* and *j* belong to the different study sites, *kr*_*i,j*_ = 0 otherwise; *kl_laussat*_*i,j*_ = 1 if the two populations *i* and *j* belong to different microhabitats within the study site of Laussat, *kl_laussat*_*i,j*_ = 0 otherwise; *kl_regina*_*i,j*_ = 1 if the two populations *i* and *j* belong to different microhabitats within the study site of Régina, *kl_regina*_*i,j*_ = 0 otherwise.

*θr*_*m*_ and *θl*_*m*_ are locus-specific parameters describing the departure from the global means of differentiation at regional (i.e. between sites) and local (i.e. between microhabitats) scales. Posteriors of *θr*_*m*_ and *θl*_*m*_ are normally distributed allowing thus to attribute an empirical Bayesian p-value (eBP-value) to each SNP. SNPs with an eBP < 1% were considered as outliers under divergent (upper tail) or homogenizing (lower tail) selection between sites (*θr*_*m*_) or microhabitat within sites (*θl*_*m*_).

Bayesian Model specifications are described in **File S3**.

For comparison, two additional methods for divergence outlier detection were also applied to our data: BayPass (Gautier, 2015) and G2D (Nielsen et al., 2009). The detailed methods associated to each analysis, as well as the results, are reported in **Files S4 and S5**, respectively. The values of the *XtX* and of the G2D statistics (respectively from the BayPass and the G2D method) from different sets of SNPs (neutral *versus* outlier, at each geographic scale) detected through our custom Bayesian framework were compared by a non-parametric Wilcoxon rank sum test with R (function ‘wilcox.text’). It has to be noted that the G2D method also required the identification of the most-likely historical demography model for the sampled populations, thus providing additional characterization of the sample.

### Phenotypic divergence in reciprocal transplants

Reciprocal transplants are ideal to test whether genetic divergence is associated with phenotypic divergence *in situ*, i.e. in real although uncontrolled conditions. A large reciprocal transplant experiment across sites and microhabitats was set-up in April-May 2011 to partition genetic and environmental sources of phenotypic differentiation in the two study sites of Laussat and Régina. The methodology is described in **File S6**.

## Results

### Genomic divergence

Bioinformatics results of *de novo* assembly and annotation are presented in **File S7**. Almost one hundred thousand (97,062) bi-allelic SNPs were identified in 25,803 contigs and 23,276 scaffolds, according to a stringent procedure, for a SNP density of 0.003/base (*sd*=0.004) with a Ti/Tv=1.80, **Table S2** and **Fig. S4**. Normalized molecular diversity π_L_ varied between 9 × 10^−6^ and 8 × 10^−3^ (**Table S3**), with little variation among populations. Among all SNPs detected, 41,050 (42.3%) were located in scaffolds with at least one predicted gene, and 56,012 SNPs (57.7%) in scaffolds without any predicted gene (total sequence length of the two scaffold types: 97828840 and 167106160 bp respectively. The average scaffold length is 3292 bp for gene-containing scaffolds and 687 bp for scaffolds without genes, and the difference between means is significant: Wilcox rank sum test, W = 6.586e+9, p-value < 2.2e-16). At last, 26,435 SNPs (27.2%) lie within a predicted gene, of which 10,863 (11.2%) within exons, while 14,615 SNPs were close to a predicted gene with a mean distance to a predicted gene of 737 bp (with min = 1bp and max = 27.2 kb).

The Bayesian model revealed low genome-wide differentiation between populations and significant departures from neutral expectations for a small subset of SNPs. Genome-wide genetic differentiation (average *G*_*ST*_ across loci) ranged between 0.026 and 0.032 at both geographical scales (credible intervals are shown in **Fig. S5**). Accordingly, genome-wide parameters of divergence estimated through *G*_*ST*_ partitioning were not higher between regions than between microhabitats within regions, as confirmed by overlapping credible intervals, **Table S4**. Coalescent modelling (**File S5**) suggested that the divergence between sites is about twice as old as divergence between sub-populations within sites. Additionally, it showed that populations at both sites globally expanded, although bottomland populations underwent expansion while hilltop populations underwent contraction. It finally provided fine estimates of migration rates between the studied populations that were in the order of 1% between sites and 20-30% between microhabitats within site.

Locus-specific departures from genome-wide divergence (i.e. outlier SNPs) were identified from the distribution of locus-specific parameters (*θr*_*m*_ and *θl*_*m*_). Between 375 and 2,045 outlier SNPs were detected to be under selection with an eBP threshold of 1%: 375 SNPs under homogenizing selection at regional scale, 2,045 SNPs under divergent selection at regional scale, 510 SNPs under homogenizing selection at local scale and 1,770 SNPs under divergent selection at local scale. After multiple-testing correction, the number of significant SNPs fell to 475 outlier SNPs (0.49% of all SNPs screened): 290 SNPs (in 268 contigs) under divergent selection at the regional scale and 185 SNPs (in 168 contigs) under divergent selection at the local scale, with no outlier shared between the regional and the local scales. Outlier SNPs under divergent selection were characterized by clines in allele frequencies between regional locations and/or local microhabitats. Clines in allele frequency across sites and microhabitats are illustrated in **Fig. 4** for different kinds of neutral and outlier SNPs.

**Fig. 3.**
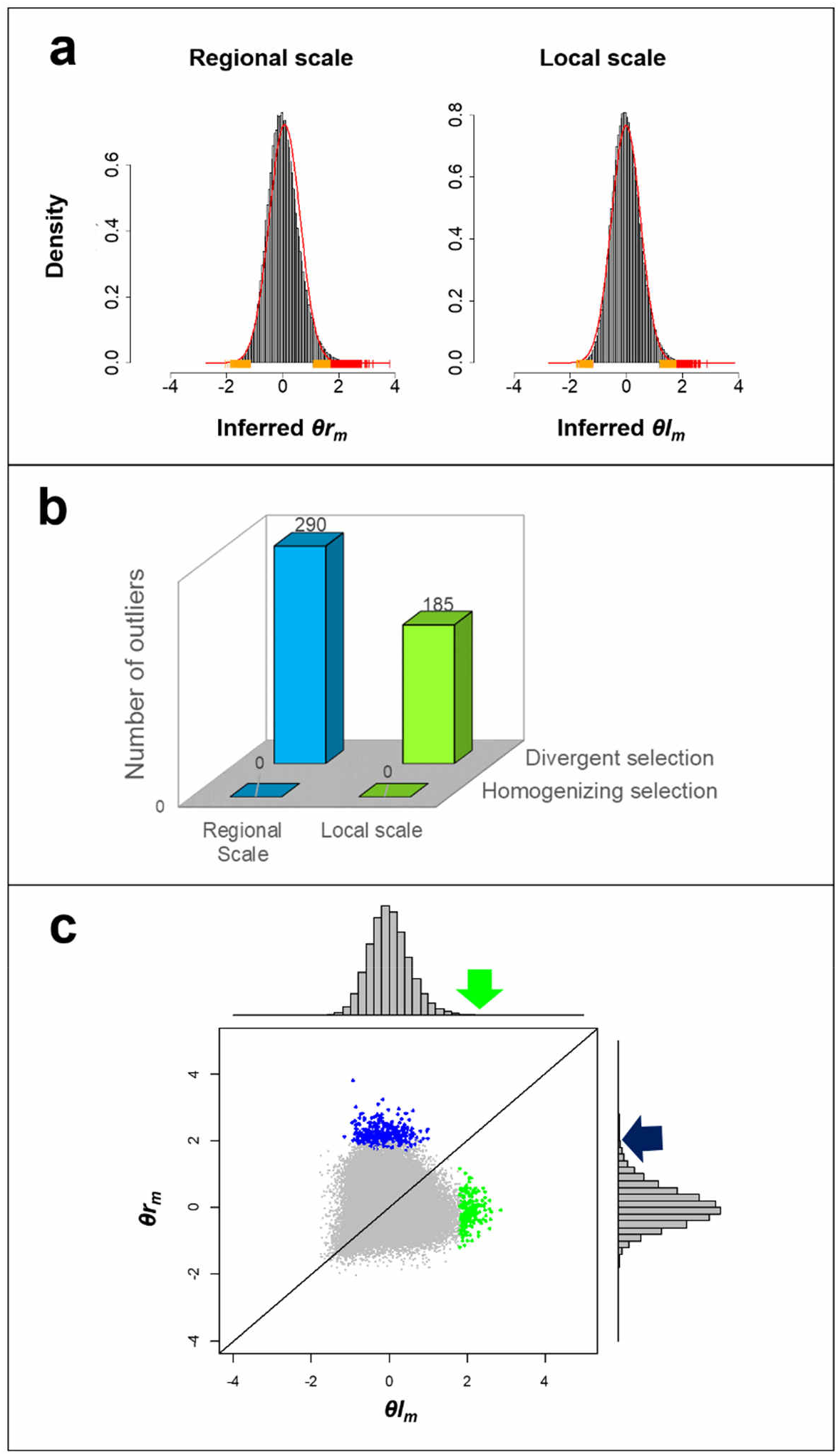
(a) Locus-specific parameters (*θr*_*m*_ and *θl*_*m*_) and normal distribution inferred by the Bayesian model. Vertical bars indicate the location of outliers under homogenizing (lower tail) and divergent (upper tail) selection. Left: regional scale (*θr*_*m*_), right: local scale (*θl*_*m*_). (b) Proportion of outlier SNPs under selection at regional (i.e. between study sites) and at local (i.e. between microhabitats within sites) scales. (c) Joint distribution of regional and local-scale locus-specific parameters inferred through the Bayesian model. Blue and green dots indicate outliers under divergent selection at the regional and local scale, respectively.

**Fig. 4.**
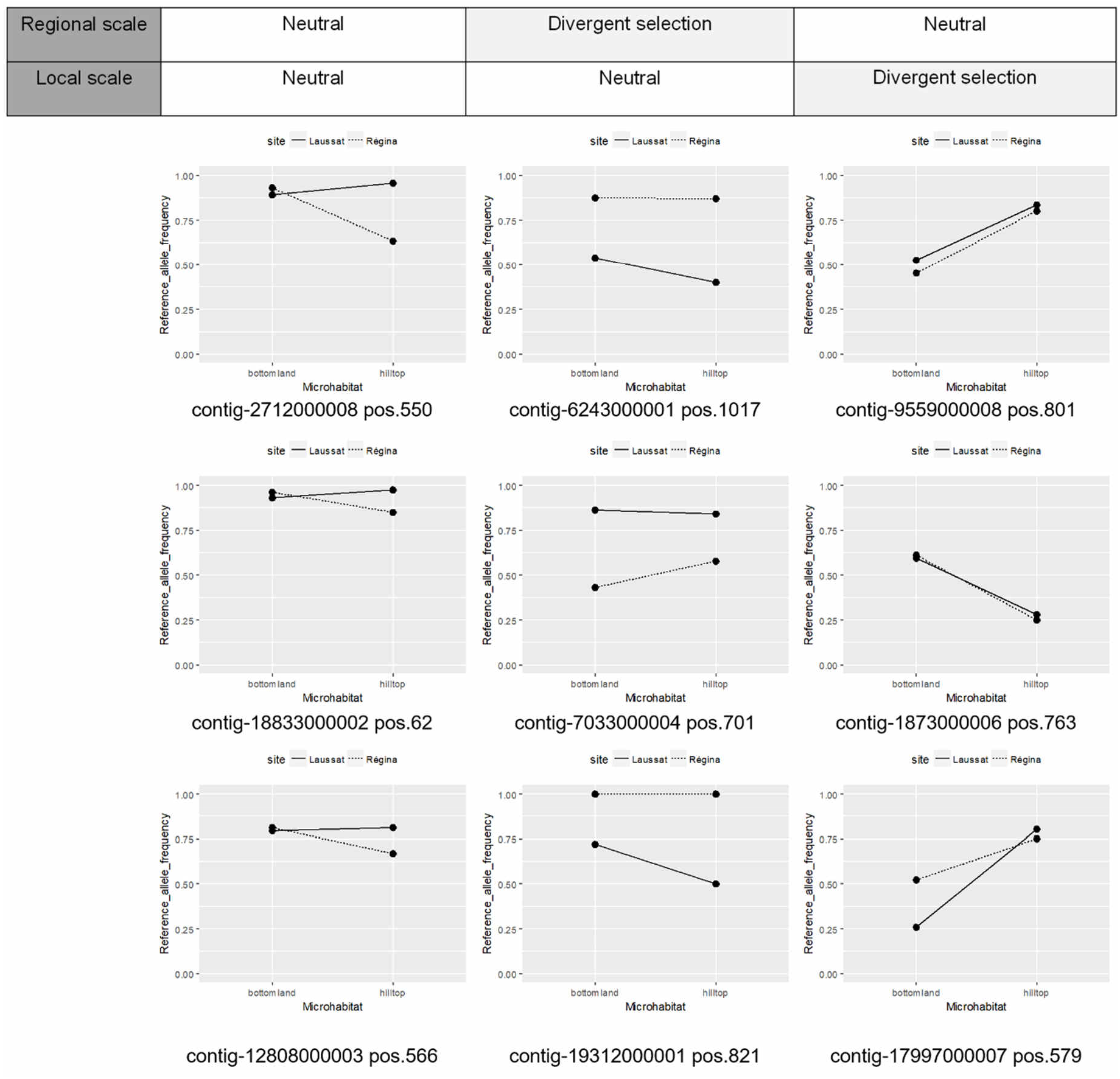
Clines in allele frequency between regional locations (i.e. study sites) and microhabitats for the different kinds of (randomly sampled) neutral and outlier SNPs at different geographical scales.

The results of the BayPass and G2D analyses are reported in **Files S4** and **S5**, respectively, providing support to the findings of our custom hierarchical model. While only six contigs showed up as outliers at both Laussat and Régina sites by the G2D, the value of the G2D statistic was significantly higher for the contigs containing outliers than for the remaining loci (**File S5**). Similarly, comparing the distribution of locus-specific *XtX* between the different sets of SNPs revealed that SNPs identified as outliers by our custom Bayesian model neutral were significantly more divergent than neutral SNPs at both regional and microgeographic scales (W statistics = 27866000 and 14542000 respectively, with p-values < 2.2e-16), **Fig. 5**. *BayPass* detected 283 outlier SNPs at the regional scale under a significance threshold of 99%, 131 of them corresponding to SNPs detected as outliers through our custom Bayesian approach (**File S4**). No microgeographic scale outliers were found by BayPass.

**Fig. 5.**
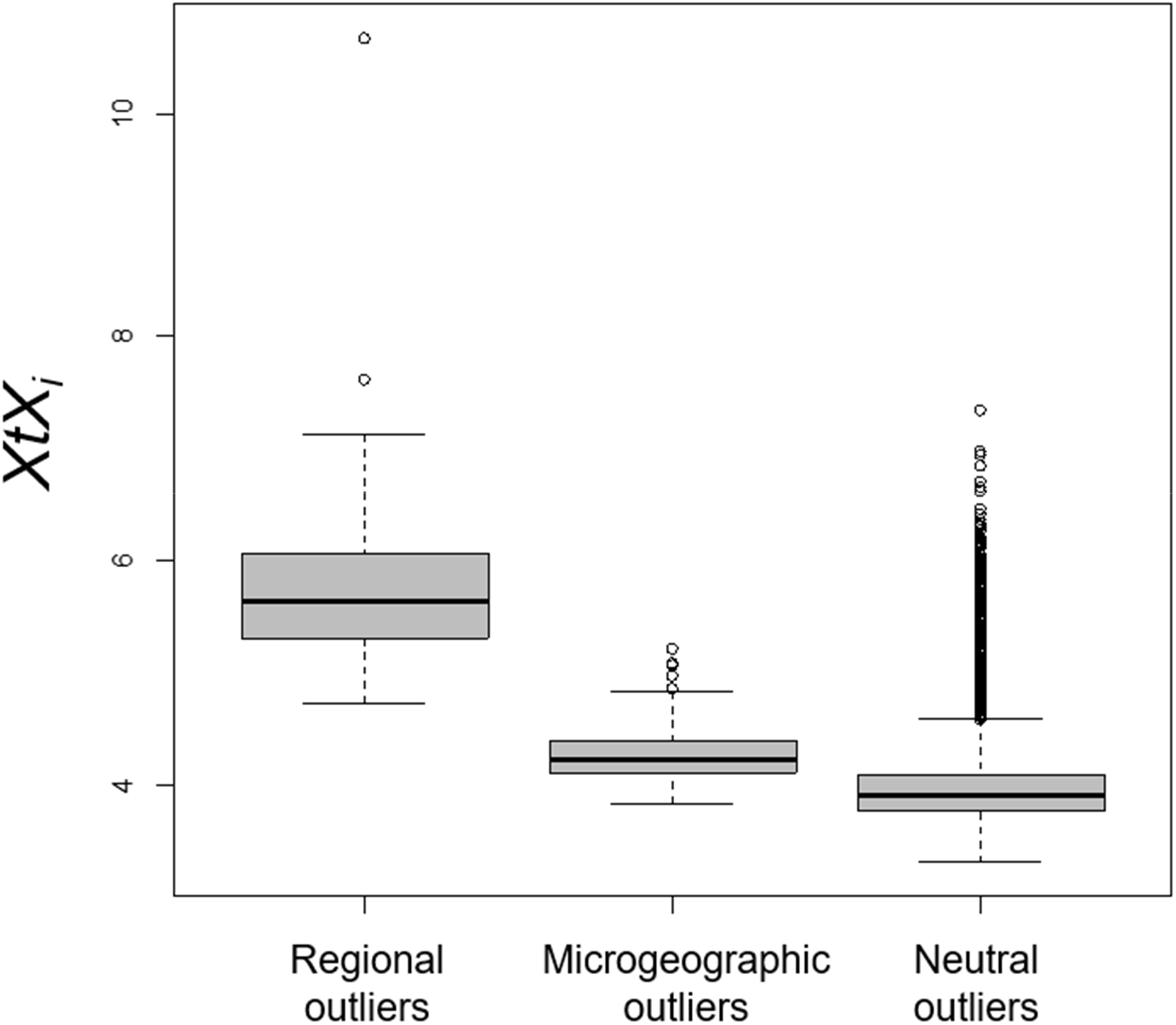
Comparison of the *XtXi* statistics estimated using BayPass across sets of SNPs: those detected as outliers at regional and microgeographic scales, respectively, and those detected as neutral by our custom Bayesian modelling framework.

A total of 140 outlier SNPs (29.5% out of 475 outlier SNPs) detected through our custom Bayesian framework were located within predicted genes, among which 39 (8.2%) were within exons, and 86 are in the neighborhood (< 5kb) of one or several predicted genes. However, outlier SNPs were not more likely to be located within exons (*X*^*2*^ = 3.9368, df = 1, p-value = 0.05), within predicted genes (*X*^*2*^ = 1.0846, df = 1, p-value = 0.30), or in the neighborhood of predicted genes (*X*^*2*^ = 3.1964, df = 1, p-value = 0.07), compared to any SNP in the entire dataset. Nevertheless, 87 GO-terms were significantly enriched in the subset of genes surrounding outlier SNPs (84 genes) compared to all predicted genes of the entire reference at the regional scale, and 100 GO-terms were significantly enriched in the subset of genes surrounding outlier SNPs (106 genes) at the local scale, **Table S5**, **Files S8 and S9**.

### Phenotypic divergence in reciprocal transplants

Patterns of genomic divergence between sites and microhabitats co-occurred with phenotypic divergence between populations occupying different regions and microhabitats. A significant provenance effect was detected for fourteen traits at the regional scale, and for ten traits at microgeographic scale, **Table 1**, **File S10** and **Fig. S7**.

**Table 1.**
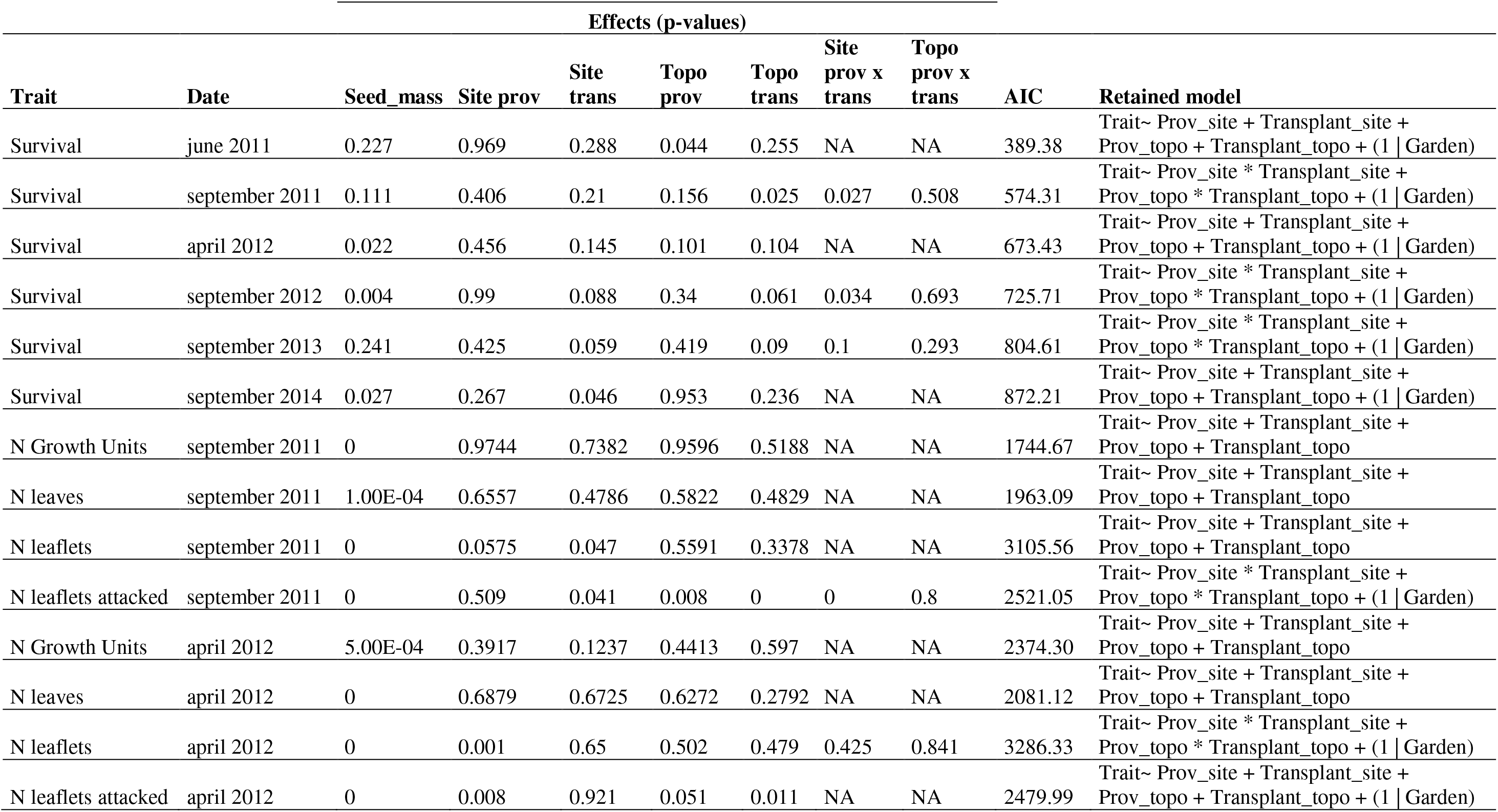
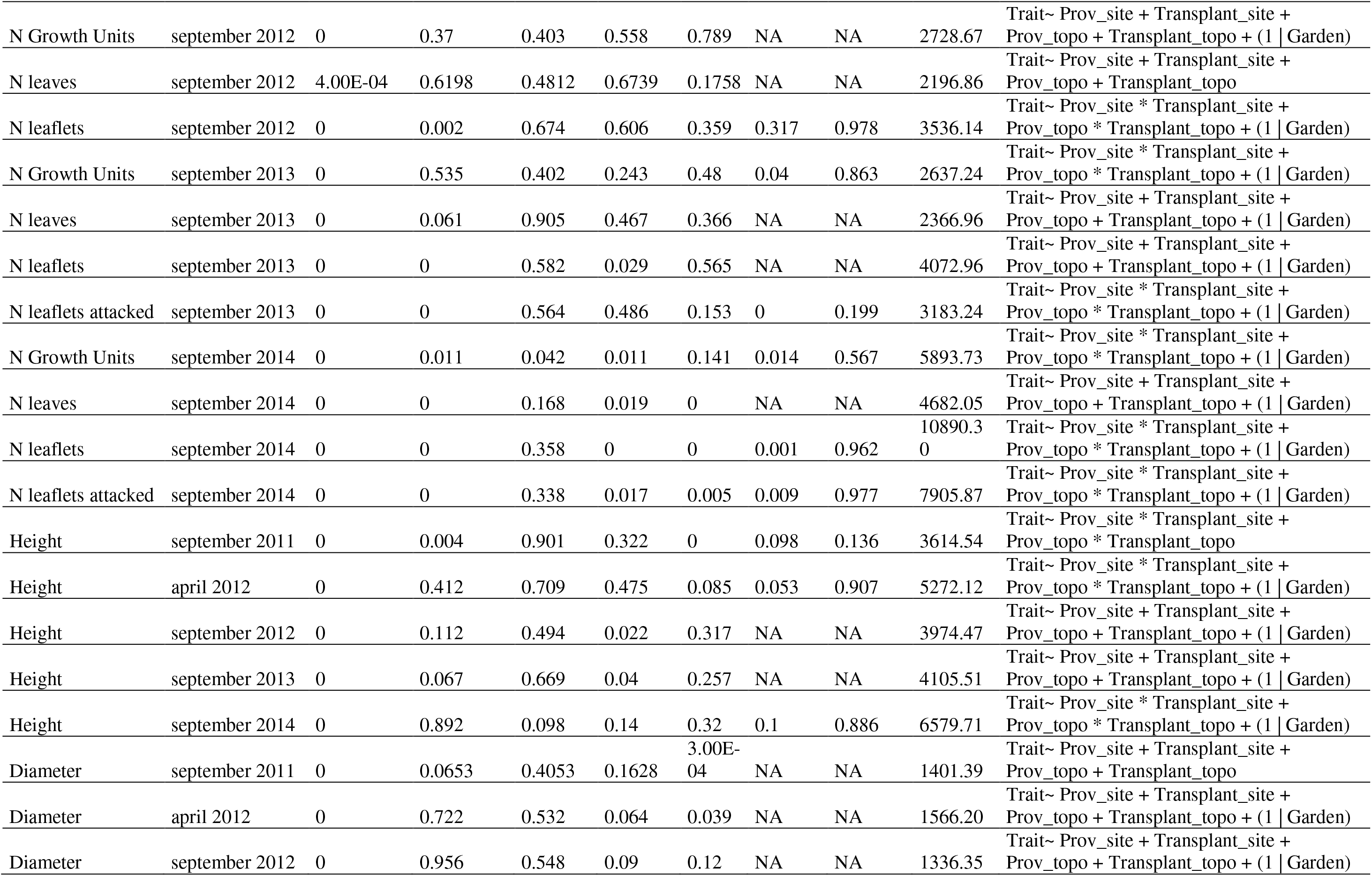
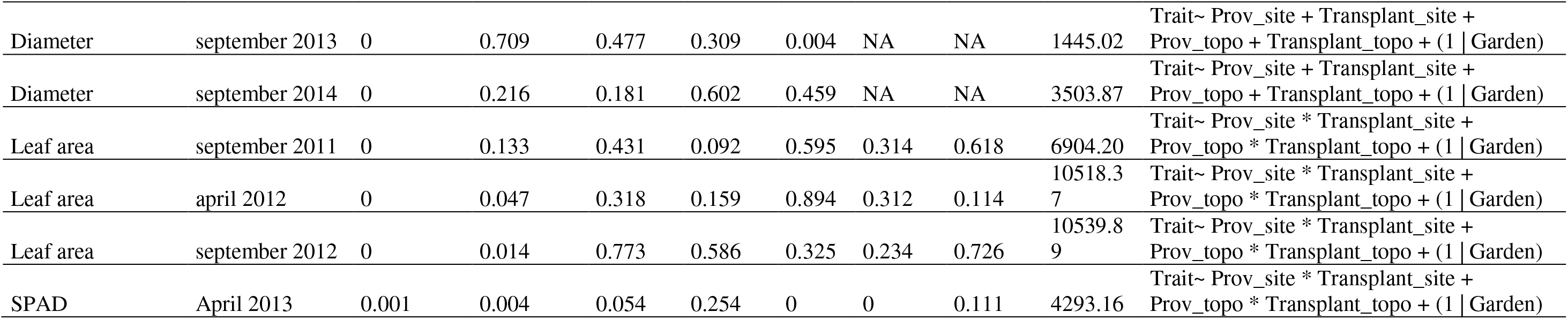
Results of reciprocal transplants - linear models of phenotypic value decomposition.

Seedlings originating from the study site of Laussat grew slowly in height but developed wider stems in diameter at early stages (in 2011), but these differences became non-significant after 2012. They however developed significantly more growth units, a trait which became significant in 2014. Leaf properties also varied between regional provenances: seedlings from Laussat established significantly more leaves (significant in 2014) and leaflets (significant in 2012, 2013 and 2014); these traits being associated with a significantly higher leaf area in September 2012. They also displayed lower chlorophyll content and more predation in 2013 and 2014 although they were less attacked by predators in 2012.

The microhabitat of provenance was also significant for several traits: seedlings from hilltops survived better in 2011 but this result became non-significant at later stages. They also grew slowly in both height (September 2012 and 2013) and diameter (September 2012) but were significantly less attacked than seedlings originating from bottomlands (2011, 2012 and 2014). Although significant in 2014, the microhabitat of provenance effect was not very pronounced regarding the number of growth units, leaves and leaflets.

Significant environmental effects of the transplant site and microhabitat were also detected. Fewer traits (four) were significant at the regional scale than at the micro-geographic scale (twelve), thus indicating a stronger influence of microenvironmental variations on seedling growth. Seedling transplanted in Laussat survived better and established more growth units (significant in 2014) and more leaflets that were also more attacked (significant in 2011). At the microgeographic scale, seedlings transplanted in bottomlands less survived (September 2011) but grew faster in both height (September 2011 and April 2012) and diameter (September 2011, April 2012 and September 2013) at early stages. They also established less leaves and leaflets (significant in 2014) that were less attacked (2011, 2012 and 2014) and had a lower chlorophyll content.

Significant Provenance-by-Environment interactions were detected providing additional evidence of local adaptation at the regional scale. A significant ‘local advantage’ in Laussat was detected on seedlings’ survival at early stages (September 2011 and September 2012), on the development of growth units (2013 and 2014), on the number of leaflets (September 2014) and on chlorophyll content. These interactions were accompanied by a ‘home advantage’ for seedlings originating from Laussat regarding the number of leaflets, the chlorophyll content, the number of growth units in 2014, and for both provenances regarding the number of growth units in 2013 and survival. Predation measurements, for their part, revealed a local disadvantage in Laussat associated with a home disadvantage for both provenances in 2011 and for seedlings originating from Laussat only in 2013 and 2014.

## Discussion

This study analyzed patterns of genomic and phenotypic divergence in wild and undisturbed forest stands of the Amazonian hyperdominant tree *Eperua falcata* in French Guiana. Previous studies (Audigeos et al., 2013; Brousseau et al., 2013, 2015) had already reported signatures of microgeographic divergence both in traits and in molecular data; nevertheless, they were not able to exhaustively investigate its evolutionary drivers, because they were based on datasets restricted to few (e.g., candidate gene approach, Audigeos et al., 2013) or anonymous markers (AFLPs, Brousseau et al., 2015), and the study of microgeographic adaptation lacked a comprehensive survey including both genomic and phenotypic levels. Here, we provide new evidence of microgeographic adaptation associated with (topographic and edaphic) habitat patchiness in this species, by identifying hundreds of SNP loci that display much higher levels of divergence at the microgeographic and regional scale than expected by pure chance, and by identifying sets of phenotypic traits that significantly diverge between populations for each geographic scale.

### Patterns of genome-wide divergence

Genome-wide (supposedly neutral) differentiation was low at both the local and regional geographic scales, ranging from 2 to 3% (**Fig. S5**), similar to what was previously found by Brousseau et al. (2015), who reported *F*_*ST*_ values ranging from 2 and 3% between microhabitats within sites, and about 3% between these study sites (based on a screening of about 1,100 AFLPs). In two additional study sites in French Guiana (Paracou and Nouragues), another study (Audigeos et al., 2013) provided comparable estimates: *F*_*ST*_ ≅ 1% between study sites and *F*_*ST*_ from 1 to 2% between microhabitats within sites. The SNP density that we estimated here (a mean of 3 SNP every 1,000 bp, **Table S2**) was quite low: Neale and Kremer (2011) for example reported that SNP frequency in trees is generally in the range 1 SNP per 100 bp. However, in our study the stringent SNP filtering probably filtered out many actual polymorphisms, leading to an under-estimation of genome-wide SNP density.

Low genome-wide differentiation suggests that high levels of gene flow in *E. falcata* regularly homogenize (neutral) genome-wide genetic diversity both between regions and between microhabitats within regions. Even though in absolute terms gene flow across sites may be relatively small, as suggested by coalescent modeling, rare long-distance seed dispersal is a common property of tree populations, and appears to have a disproportionate effect on genetic connectivity, potentially even facilitating local adaptation (Kremer et al., 2012). In general, the amount of divergence observed here is comparable to what has been found in more intensively studied temperate or tropical tree species (e.g., Alberto et al., 2010; Dick & Heuertz, 2008; Scalfi et al., 2014). The consequences of recurring inter-habitat gene flow on population adaptability could be paradoxical. On one hand, extensive gene flow can allow the spread of advantageous alleles between populations, thus granting high adaptive potential for populations established in heterogeneous environments. On the other hand, gene flow may also counteract the effect of natural selection, with migration load halting adaptive divergence of populations into locally adapted subpopulations (De Kort, Vandepitte, Mergeay, Mijnsbrugge, & Honnay, 2015; Le Corre & Kremer, 2012; Lenormand, 2002; Pujol et al., 2018). The deleterious effect of gene flow on adaptive divergence could explain the low proportion of outliers detected (see below); another explanation may be the highly polygenic nature of adaptive traits, which would prevent individual underlying genes to diverge in frequency (Le Corre & Kremer, 2012). Nevertheless, high gene flow (and resulting low genome-wide differentiation) did not completely swamp the signal of adaptive divergence between populations.

### Local adaptation across geographic scales

Locus-specific (putatively adaptive) departures from genome-wide divergence were detected for 0.49% of the 97,062 SNPs analyzed. Results at the single SNP level were confirmed by trends at the contig level, showing similar trends of microgeographic differentiation. Moreover, a comprehensive reciprocal transplant experiment also corroborated evidence of local adaptation by revealing phenotypic differences at both regional and microgeographic scales in seedlings at early developmental stages, when natural selection is expected to be stronger (Petit & Hampe, 2006).

Significantly more outliers were detected at the regional than at the microgeographic scale: 290 *versus* 185 outlier SNPs (*X*^*2*^ = 22.826, df = 1, p = 1.77 × 10^−6^). This result can be explained as the logical consequence of an order of magnitude difference in the geographical distances between populations: 300 meters between microhabitats within sites versus 300 km between study sites. Indeed, ecological conditions are probably more variable, and divergent selection stronger, at the regional scale (mainly driven by climatic differences associated with different rainfall regimes) than divergent selective pressures between microhabitats separated by hundreds of meters (driven by differences in soil properties), and the effect of divergence would be reinforced by more limited gene flow. The proportion of outliers detected over microgeographic scales, although smaller, is nevertheless relevant. Because the Bayesian model focuses on replicated footprints of adaptation only, it is extremely unlikely that the detected outliers could have been structured between microhabitats in two different regions independently purely by chance (Savolainen, Lascoux, & Merila, 2013). Moreover, no outlier was detected jointly at both regional and microgeographic spatial scales, as shown by the heart-shaped joint distribution of regional- and local-scale locus-specific parameters (**Fig. 3**). Instead, outliers were always specific to only one spatial scale, suggesting that different portions of the genome respond to different types of environmental constraints. Reciprocal transplants also indicated that different sets of traits diverge between populations at regional and microgeographic scales. Although it is impossible to link the analyzed phenotypic traits to one or several SNPs detected without a comprehensive association study, these results suggest that different environmental drivers would be at stake and that, in future studies, local adaptation in wild populations should be investigated by integrating different spatial scales, including microgeographic habitat patchiness, into the sampling design.

### Methodological considerations

Whole-genome sequencing of population pools has opened a new window of opportunity, especially for non-model species. Their analysis however remains complicated as it raises many methodological and statistical challenges, even more in the case of complex – presently *replicated* – sampling designs.

The Bayesian modelling framework implemented here was specifically designed to detect replicated footprints of divergent selection between microhabitats under an explicit hierarchical model of neutral divergence both between and within study sites, thus taking into account variable levels of gene flow across spatial scales. This custom framework however suffers from several limitations, including computation time, estimation uncertainty and detection power.

Computation time limitation was overcame by splitting the SNP dataset into ten independent subsets. The fact that neutral parameters were similar across subsets suggests that locus-specific parameters were estimated under similar conditions of hierarchical neutral divergence between populations across subsets (**Fig. S6**). This is a natural consequence of the largely fragmented reference assembled *de novo*: SNPs were randomly scattered across the genome and they can therefore be considered as independent. Although SNPs located on the same scaffolds are physically linked, they did not cause major discrepancies in the estimation of neutral parameters across subsets.

Assessing the chances of *F*_*ST*_-based selection tests to detect false-positives and to miss true-positive outliers is another challenge, and this issue strongly depends on the size of the dataset. While the power of our custom Bayesian model was not estimated for very large numbers of loci by Brousseau et al. (2016), we included in the preset study an additional correction for multiple-testing (with a FDR threshold of 10%) to deal with the large number of SNPs analyzed. Although linking directly FDR with FNDR, and subsequently with the power of our model is difficult, there exist theoretical and numerical expectations of the False Non-Discovery Rates (FNDR) of multiple tests based on the chosen FDR threshold. In particular, Genovese & Wasserman (2002) show that, for an FDR threshold of 5%, FNDR is always below 20% for cases where most tests are expected to be true negatives. We can expect that only a small minority of loci is under divergent selection at our small phylogenetic and spatial scales. Finally, Sarkar (2006) defines the power π_0_ = 1 – FDR – FNDR; therefore, in our case, power should be in the order of 1− 0.1 - O(0.2), that is, no larger than 70%, and probably somewhat smaller.

Another source of bias lies in the inference of population-specific allele frequencies and genetic differentiation from read counts in population pools because of the double sampling uncertainties, (1) when sampling individuals from a population, and (2) when sampling reads from the pool of haploid sequences during sequencing (Ferretti, Ramos-Onsins, & Pérez-Enciso, 2013; Hivert, Leblois, Petit, Gautier, & Vitalis, 2018). This limit was tackled by applying a library-level coverage threshold to detect SNPs: a threshold of 20X was used, that is equal the number of diploid individuals (or half the haploid sample size) in each population pool. Applying library-level coverage threshold to infer library-specific allele frequency is mandatory to reduce the probability of missing alleles at low frequency, although fluctuations in the sampling process are unavoidable and sequencing all distinct lineages in a pool of 20 diploid individuals remains unlikely even at 40X coverage (Ferretti et al., 2013). Aware of these limitations, uncertainties in allele frequencies were however further integrated in estimates of pairwise genetic differentiation (*G*_*ST*_) between libraries under the Bayesian framework.

It is also important to stress that our Bayesian approach may miss footprints of polygenic adaptation, if adaptation acts in different ways in the two study sites (through different selection strengths and/or different loci affected) which is known to be common in wild trees (Berg & Coop, 2014; Harrisson et al., 2017; Lind et al., 2017). Even if the Bayesian model employed here proved its robustness to detect footprints of polygenic adaptation caused by weak selection strengths (*s*=0.05) from genotype data in simulations (Brousseau et al., 2016), it may still miss footprints of polygenic adaptation if the strength of selection is variable among loci (Sork et al., 2013). Indeed, *F*_*ST*_-based methods are more prone to detect adaptation caused by loci of large effects than by loci with small effects, especially since adaptation occurs with ongoing gene flow (Hedrick, 2006; Le Corre & Kremer, 2012; Savolainen et al., 2013; Yeaman & Whitlock, 2011). In other words, the set of outliers detected is probably enriched with loci of large effects. Moreover, it is possible that the genetic architecture of traits selected at different geographical scales differs, thus explaining both the differences in numbers of outliers detected and lack of overlap. Indeed, the reciprocal transplant experiment showed that different sets of traits inherently diverged between sites and between microhabitats within sites. It is also likely that traits closer to metabolic pathways (such as chlorophyll content, possibly chemical defense against grazers) are less polygenic than complex growth traits, and that therefore the chances of identifying single-locus outliers vary between these different sets of traits.

Beyond theoretical considerations, the results obtained were further compared with two other widespread selection tests suitable for analyzing Pool-Seq data, G2D and *BayPass*, respectively. Applying different methods often produce partially different results, because of inherent conceptual and mathematical differences in methods formulation and implementation. For example, G2D can detect replicated footprints of selection taking into account the demographic history of the studied populations at the contig-level, not at the SNP level. *BayPass v2.1*, for its part, takes into account complex neutral structuring through the inference of a genome-wide variance-covariance matrix and allows detecting SNPs that are over-differentiated between standing populations and the ancestral one, without however distinguishing outliers caused by crossed-factors nor repeated binary contrasts. The identity and quantity of outlier contigs or SNPs detected in the present study thus vary between these different selection tests. Comparing the distribution of locus-specific differentiation (G-values and *XtX*) between the different sets of SNPs detected by our custom Bayesian framework however revealed a shared trend: the extent of differentiation was always higher for outlier SNPs than for neutral SNPs, indicating that the sets of SNPs detected as outliers at both regional and microgeographic scales were enriched in SNPs influenced by divergent selection although they possibly hide false-positive outliers.

### Microgeographic adaptation

This study was primarily designed to test the hypothesis of replicated microgeographic adaptation within two independent study sites, and the regional scale was thus taken into account to avoid confounding footprints of adaptation between large-scale and small-scale factors, and to compare the strength of adaptive divergence between spatial scales (in terms of number of SNPs affected). For these reasons, our interpretation thus focuses on footprints of microgeographic adaptation only thereafter.

Although detecting adaptive divergence between regions at geographic scales on the order of hundreds kilometers is not surprising, detecting patterns of microgeographic adaptive divergence on the order of hundreds meters is less common in tree populations. By providing evidence of adaptation to microgeographic habitat patchiness at both genomic and phenotypic levels, our results agree with previous studies showing soil variation as a driver of adaptation in trees and, more specifically, in the Neotropics (Audigeos et al., 2013; Brousseau et al., 2015).

Topography-linked microgeographic patchiness is likely driven by a complex interaction of abiotic and biotic factors (including soil types (Misiewicz & Fine, 2014), water-logging and flooding events, water availability (Clark, Palmer, & Clark, 1999), seasonal soil drought (Markewitz, Devine, Davidson, Brando, & Nepstad, 2010), soil fertility (John et al., 2007), light availability (Ferry, Morneau, Bontemps, Blanc, & Freycon, 2010), competition and predation (Fine et al., 2004)). At the community level, the study by Fortunel et al. (2016) already suggested a mechanism through which microgeographic adaptation may take place: seedling mortality of *terra firme* species was high when planted in seasonally flooded habitats. At the intra-specific level, previous studies have reported examples of adaptation to soil nutrient composition such as potassium deficiency in *Cornus florida* (Cornaceae) (Pais, Whetten, & Xiang, 2017), to aluminum and phosphate ion concentrations in *Pinus contorta* (Pinaceae) (Eckert, Shahi, Datwyler, & Neale, 2012), to substrate age in *Metrosideros polymorpha* (Myrtaceae) (Izuno et al., 2017), to soil types in *Abies religiosa* (Pinaceae) (Méndez-González, Jardón-Barbolla, & Jaramillo-Correa, 2017), or to soil water availability in *Pinus albicaulis* (Pinaceae) (Lind et al., 2017). Preliminary evidence of microgeographic adaptation has been recently reported in Neotropical tree species, including *E. falcata* (Audigeos et al., 2013; Brousseau et al., 2013, 2015; Torroba-Balmori et al., 2017). We provide here additional evidence of microgeographic adaptation in *E. falcata* at the genomic scale, also corroborated by significant effects of the microhabitat of provenance at the phenotypic level.

It is, however, difficult to identify the exact genomic targets of natural selection, particularly in non-model species. First, because linking complex phenotypes (growth and leaf traits) with the SNPs analyzed is impossible without an accurate association genomics approach. Second, because of potential hitchhiking between causal mutations and the analyzed SNPs, and because of the lack of chromosome-level genome assembly as well as no exhaustive genome-wide estimates of linkage disequilibrium in this species (but linkage disequilibrium is nearly zero beyond few hundred base pairs in coding sequences, Audigeos et al., 2013). Despite these limitations, the identity of the genes neighboring outlier SNPs may provide useful information, as many ‘divergence’ outlier SNPs detected here were located within or close to (< 5 kb) protein-coding genes involved in a variety of biological functions (**Files S8 and S9**), with 106 genes neighboring a genomic footprint of microgeographic divergence (i.e. one or several outlier SNPs) and 100 Gene Ontology terms (corresponding to 49 genes) significantly enriched in the subset of genes neighboring outlier SNPs, **Table S5**. Some of them are particularly meaningful regarding their biological significance in terms of survival, growth, defense and reproduction (**Files S8 and S9**). Functional annotation and subsequent enrichment tests are however to be taken with caution. Although they can provide useful clues about the biological functions of the genes identified, they do not necessarily provide a proof of the results functional meaning (Pavlidis, Jensen, Stephan, & Stamatakis, 2012).

We detected several genes involved in plant response to stress close to SNPs detected as outliers at the microgeographic scale. For example, a glutaredoxin-C11 (predgene_010110) involved in managing the level of reactive oxygen species (ROS) in plant cells (Mittler, Vanderauwera, Gollery, & Van Breusegem, 2004), a H_2_O_2_-responsive gene encoding a metallopeptidase M24 family protein which is up-regulated under stress in *Arabidopsis thaliana* (Ordoñez et al., 2014), and a gene encoding a plant cysteine oxidase 4-like (predgene 003921) involved in response to flooding and soil hypoxia (Weits et al., 2014; White et al., 2017) were detected. Oxidative stress is very common in bottomlands, where water-logging and seasonal flooding events cause soil hypoxia and oxygen deprivation in root cells, leading to the production of ROS (Blokhina, Virolainen, & Fagerstedt, 2003). This observation is concordant with a previous candidate-gene approach in which a catalase (i.e. a gene involved in ROS detoxification) has been reported under selection in this species (Audigeos et al., 2013). We also detected five genes with possible implications in lipid biosynthesis (predgenes 006075, 019500, 002577, 001066, 000689), as for example a palmitoyl-acyl carrier protein thioesterase which is known to play an essential role for plant growth and seed development (Bonaventure, Salas, Pollard, & Ohlrogge, 2003; Dörmann, Voelker, & Ohlrogge, 2000). Lipid metabolism is of high importance for plant development, from early germination to reproduction. Indeed, lipids are the primary material for cutin, and they are largely involved in stress signaling and resistance (Okazaki & Saito, 2014). We also found two genes involved in proline amino acid biosynthesis (predgenes 007023 and 007024), which is accumulated in response to drought stress in plants (Fu, Ma, Chen, Gu, & Gong, 2018). A calmodulin-like protein (probable CML48, predgene 022002) was also detected in the neighborhood of a microgeographic outlier. Calmodulin and calmodulin-like plant calcium sensors have evolved unique functions of high importance in the regulation of plant development and plant response to stress (Perochon, Aldon, Galaud, & Ranty, 2011). We also found a probable cellulose synthase A (predgene 000745) whose mutations can alter morphology or induce resistance to cellulose biosynthesis inhibitors as reported in *A. thaliana* (Richmond, 2000), a gene for plant metal tolerance (predgene 014347), and a mRNA-decapping enzyme-like (predgene 011170) which is of importance for early seedling development (Goeres et al., 2007).

Two genes mediating biotic interactions - through plant defense against pathogens and recognition of symbiotic bacteria – were also detected as potential targets of microgeographic adaptation. In particular, plant chemical defense against herbivores and pathogens is likely to play a key role in microgeographic adaptation; previous studies have experimentally demonstrated that a trade-off between growth and antiherbivore defenses strengthens habitat specialization between congeneric species within large Neotropical plant families (e.g., Annonaceae, Burseraceae, Euphorbiaceae, Fabaceae, Malvaceae) (Fine et al., 2004, 2013, 2006). Here, we detected a chitinase (predgene 000254), a family of enzymes known for their role in resistance to pathogens (Grover, 2012; Punja & Zhang, 1993). This is fully consistent with the finding of significant provenance effects of seedling predation (estimated by the number of leaflets attacked) in the present study, which suggests biotic selective pressures exerted by herbivore communities should contribute to microgeographic adaptation. We also discovered a gene involved in recognition of symbiotic bacteria (LysM receptor-like kinase, predgene 002974) which enables the model legume *Lotus japonicus* to recognize its bacterial symbiont (Madsen et al., 2003; Radutoiu et al., 2003; Ryals et al., 2008). Symbiotic nitrogen fixation is a specific feature of legumes, and nutrient availability largely varies across microhabitats (Ferry et al., 2010). The presence of genes involved in the perception of symbiotic bacteria close to genomic footprints of microgeographic adaptation in this supposedly non-nodulating legume (Villadas et al., 2007) is particularly interesting. However, accurate functional characterizations of the proteins surrounding outlier SNPs are required to draw robust conclusions about the physiological processes involved in microgeographic adaptation.

### Microevolution in the (Neo)tropics

Although largely exploratory, our results open a new window into the evolutionary processes underlying the maintenance of genetic and functional diversity, and support the hypothesis of niche evolution as a driver of habitat specialization and diversification (Leibold, 2008). Microgeographic adaptation driven by a complex interaction of abiotic and biotic factors may be an initial step towards habitat specialization and ecological (sympatric) speciation by trees (Feder, Egan, & Nosil, 2012; Savolainen et al., 2006). While it is true that microgeographic adaptation is not at all restricted to (Neo)tropical trees (e.g., Lind et al., 2017; Scotti et al., 2016), the extreme patchiness of micro-environmental conditions and the overall levels of competition for resources and space observed in tropical ecosystems may amplify the specialization of trees to specific microhabitats. In this study we provide support for this process at the intra-populational level in an Amazonian tree species. However, the efficacy of microgeographic adaptive divergence at a small but non-negligible fraction of loci in driving ecological speciation (as opposed to the effect of genome-wide neutral divergence, caused by strong restrictions in gene flow, in assisting allopatric speciation) requires further investigation. In particular, a comprehensive genome-wide association study in our reciprocal transplant would provide detailed information on the functional role of the SNPs and genes potentially under selection.

It is also possible that microgeographic adaptation contributes to the generalist and hyperdominant distribution of *E. falcata* (ter Steege et al., 2013). In this view, local adaptation to habitat patches would maintain multiple adaptation optima (i.e. multiple genetic combinations of alleles at different loci), thus pre-adapting the populations to a multitude of environmental situations. Indeed, two evolutionary processes can theoretically allow populations to cope with environmental changes, such as global climate change, and avoid extinction (Aitken, Yeaman, Holliday, Wang, & Curtis-McLane, 2008; Harrisson et al., 2014): populations can migrate to new areas to track ecological niches, causing shifts in the species’ distribution, and/or adapt locally to the new conditions through the action of natural selection (Hoffmann & Sgro, 2011). These processes are likely to occur simultaneously and populations’ response to global climate change can be viewed as a ‘race where populations are tracking the moving optima both in space and time’ (Kremer et al., 2012). From the standpoint of the entire metapopulation of *E. falcata* in the region, local adaptation to multiple optima would be tantamount to population-level balancing selection, which is considered to be a major component of adaptive potential (Barrett & Schluter, 2008; Delph & Kelly, 2014). Such a reservoir of adaptively useful genetic diversity may help the species deal with environmental challenges.

To conclude, here we provide new, genomic-scale, evidence of divergent selection at both regional and microgeographic scales in the Amazonian hyperdominant *Eperua falcata*, suggesting that microgeographic environmental heterogeneity caused by variations in topography and edaphic factors creates the conditions for local adaptation in lowland Neotropical rainforests. We notably detected signals of adaptive divergence at many SNPs, whose surrounding genes may ultimately constitute the genomic basis for local adaptation. Accurate functional annotation further provided some interesting clues as to the biological processes targeted by divergent selection, emphasizing the potential roles of abiotic stresses related to water-logging and drought, and of biotic interactions, in driving local adaptation. We firmly believe that new advances in next-generation sequencing technologies and in Bayesian methods will contribute to fill the gaps in our understanding of trees evolution in the Neotropics, and we strongly recommend the integration of different spatial scales (including microgeographic habitat patchiness) in future studies addressing the processes of adaptation and diversification in lowland rainforests of Amazonia.

## Acknowledgements

This study has been funded by the ANR ‘*BIOADAPT FLAG’* (ref. ANR-12-ADAPT-0007-01) and benefited from an ‘*Investissement d’Avenir*’ grant (CEBA, ref. ANR-10-LABX-25-01), both managed by the French *Agence Nationale de la Recherche*. Louise Brousseau has been funded by a young investigator grant ‘*Contrat Jeune Scientifique*’ (CJS) of the *French National Institute for Agronomical Research* (INRA). We are grateful to the *Genotoul bioinformatics platform Toulouse Midi-Pyrenees* (Bioinfo Genotoul) for providing computing and storage resources. We thank Saint-Omer Cazal (INRA, UMR EcoFoG) and Julien Engel (IRD, UMR AMAP) for technical assistance and botanical identifications. We thank Bruno Ferry (AgroParisTech, UFR FAM), Hendik Davi (INRA, URFM) for sharing soil and functional data of adult tree populations. We thank Dr Anna Stavrinides for helpful discussions about the functional interpretation of the results. We are also grateful to Victoria Sork, Andrew Eckert and two anonymous reviewers for their very useful comments on this study.

## Author contribution

LB, ED and IS designed the experiment. LB set up the reciprocal transplant experiment and analyzed related phenotypic data under the supervision of IS and ED. LB sampled leaf material, extracted and conditioned DNA, implemented the bioinformatics pipeline and analyzed SNPs data with the custom Bayesian framework and *BayPass* under the supervision of IS and GVV. IS computed basic diversity statistics and implemented G2D analyses. All authors wrote the article.

## Supplementary figures and tables

**Supplementary figures S1 to S7.**

**Supplementary tables S1 to S5.**

## Online supporting information

<u>**Supplementary file S1.**</u> Soil characteristics of the two sites and on growth and functional characters of the sampled trees

<u>**Supplementary file S2.**</u> Bioinformatics pipeline

<u>**Supplementary file S3.**</u> Bayesian model specifications

<u>**Supplementary file S4.**</u> BayPass – methodology & results

<u>**Supplementary file S5.**</u> G2D – methodology & results

<u>**Supplementary file S6.**</u> Reciprocal transplants – methodology

<u>**Supplementary file S7.**</u> Bioinformatics results

<u>**Supplementary file S8.**</u> BLASTp and functional annotation of genes neighboring (< 5 kb) ‘divergence’ outlier SNPs at the microgeographic scale.

<u>**Supplementary file S9.**</u> Interactive gene network to easily navigate through the functional annotation of genes neighboring ‘divergence’ outlier SNPs (< 5 kb) at the microgeographic scale. Green bubbles indicate predicted genes; blue and pink bubbles indicate Gene ontology terms, with size depending on the number of predicted genes neighboring an outlier SNP flagged with each GO term; pink bubbles indicate GO terms significantly enriched between the subset of predicted genes neighboring outlier SNPs compared to the all predicted genes.

<u>**Supplementary file S10.**</u> Reciprocal transplants – linear models raw outputs

