## Supplementary material for "Genomic and phenotypic divergence unveil microgeographic adaptation in the Amazonian hyperdominant tree *Eperua falcata* Aubl. (Fabaceae)": Fig. S1 to S6

**Fig. S1.** *Eperua falcata* Aubl. (Fabaceae, Caesalpinioideae). (a) Tree, 8-40 m tall, trunk 20-80 cm in diameter. (b) Paripinnate leaves with falcate leaflets, as viewed from the ground. (c) Pendant inflorescence. (d) Explosive pods. (e) Hermaphrodite flowers. (f) Trunk slash. (e) Young seedling understory. (*Photos by Julien Engel).*

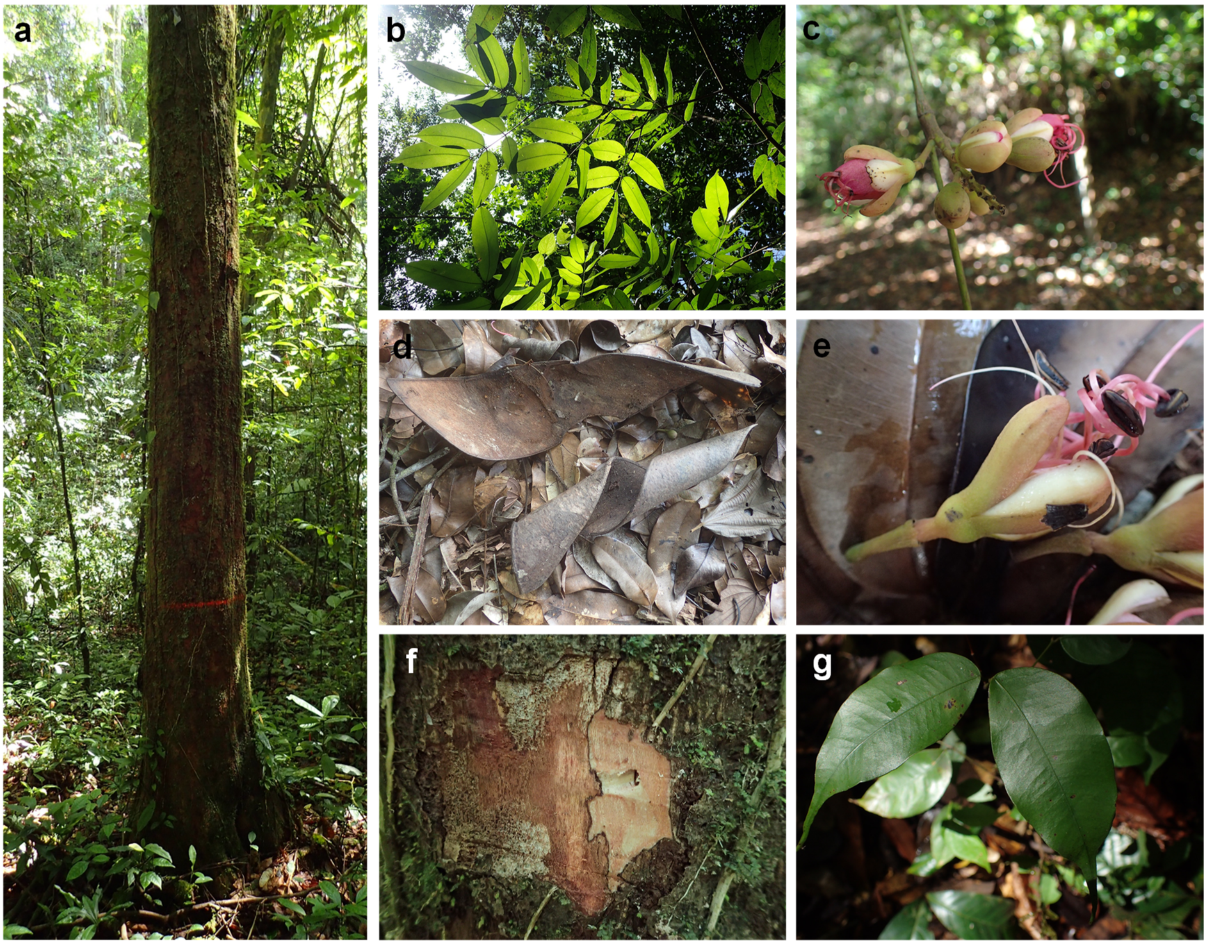

**Fig. S2.** Soil water content (%) in the two study sites and the two microhabitats within each study site in 2011 and 2012.

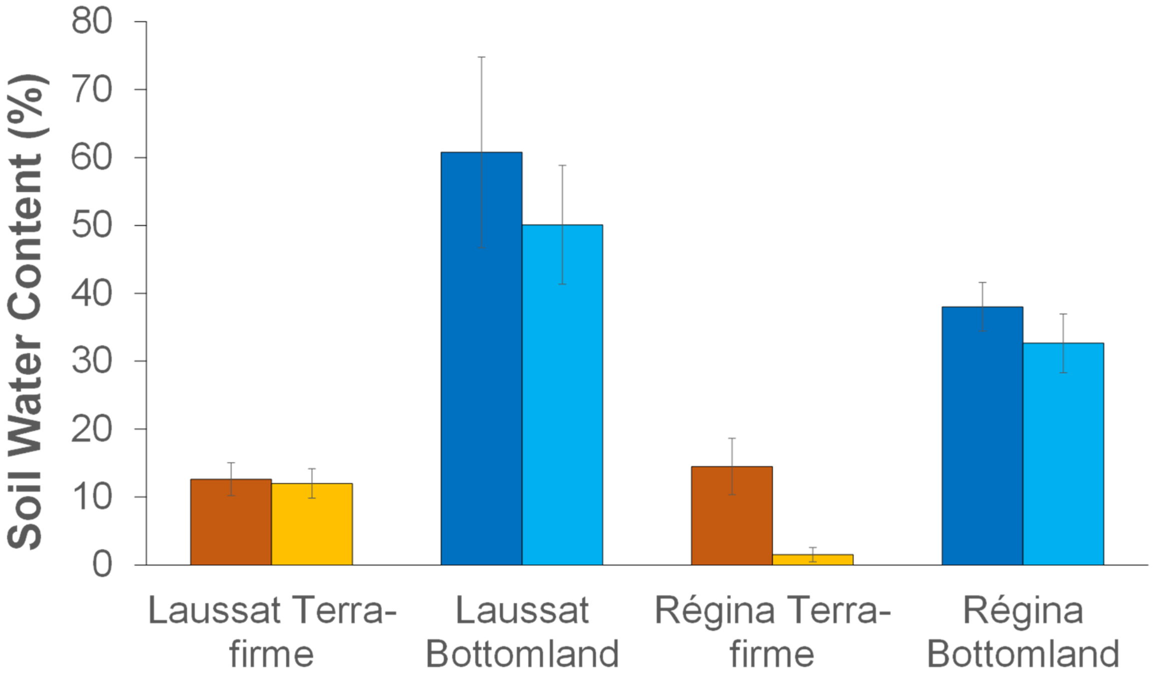

**Fig. S3.** Assembly Statistics. (A) Depth distribution (mean = 57.81X). (B) Proportion of the genome mapped by one, two, three and four libraries respectively.

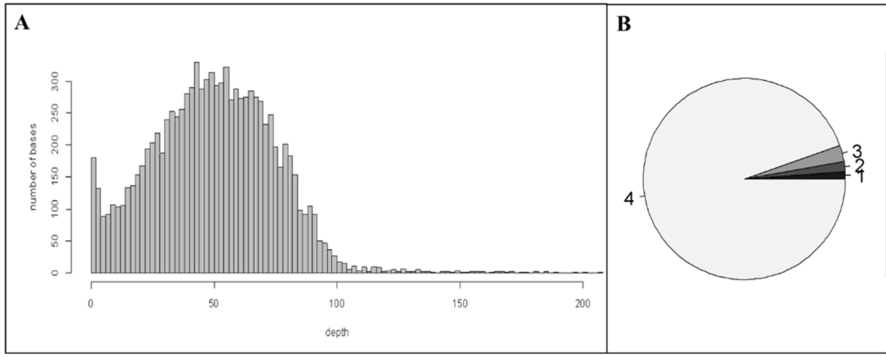


| | Proportion of assembled bases (%) |
|---|---|
| 1 lib mapped | 1.22% |
| 2 lib mapped | 1.5% |
| 3 lib mapped | 2.53% |
| 4 lib mapped | 94.75% |

**Fig. S4.** SNP Statistics: number of substitutions (transitions, transversions) and indels.

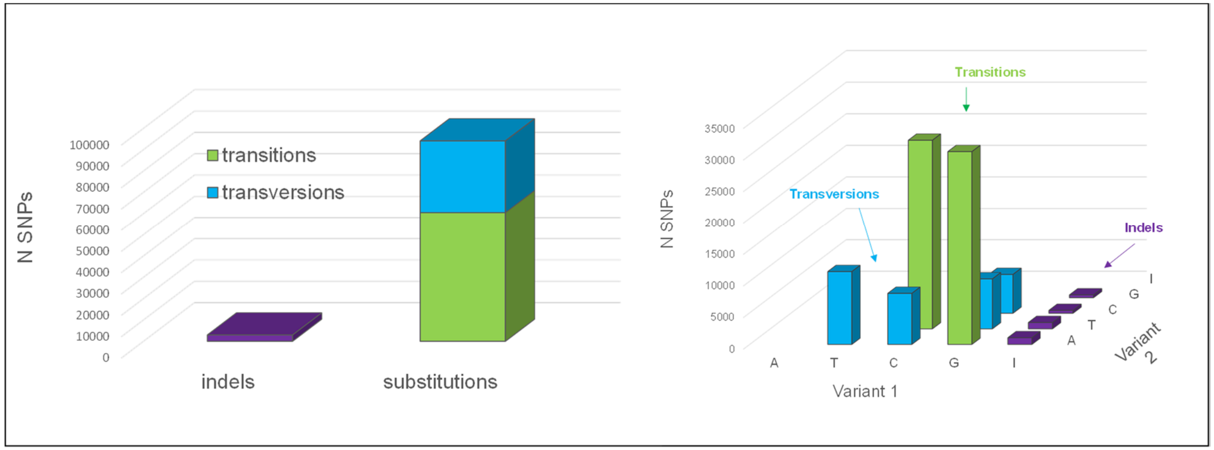

**Fig. S5.** Distribution and average $G_{ST}$ between pairs of populations inferred by the Bayesian model. R TF= Regina Terra-firme, R BL= Regina Bottomland, L TF= Laussat Terra-firme, L BL= Laussat Bottomland.

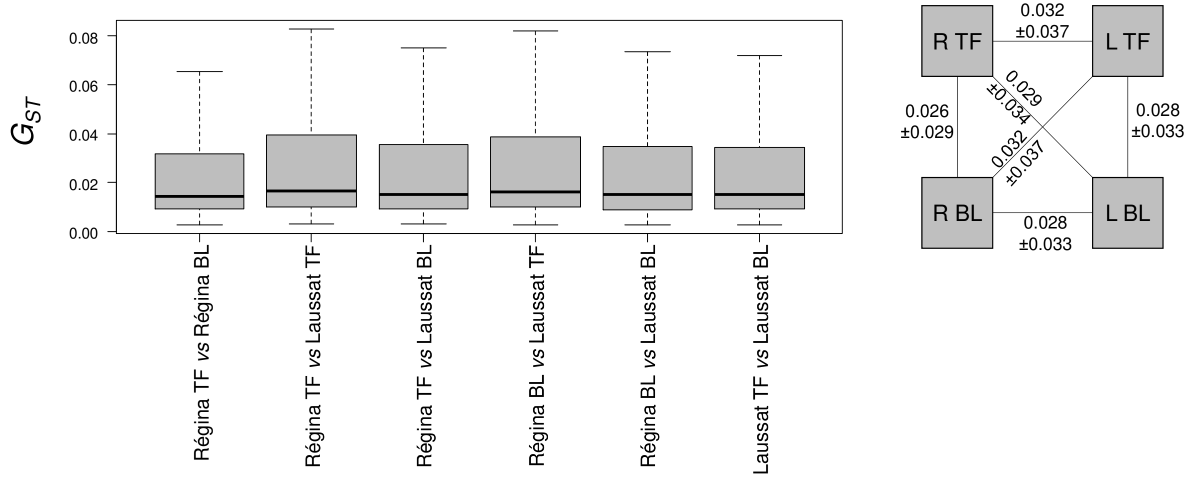


**Fig. S6.** Genome-wide parameters estimates across subsets.

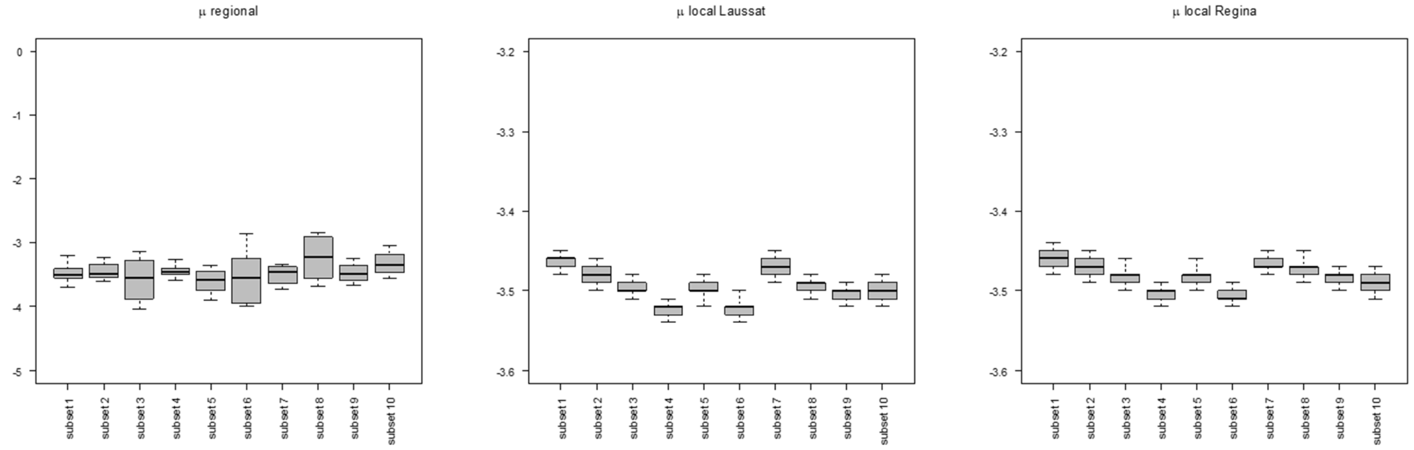
