## Supplementary material for "Genomic and phenotypic divergence unveil microgeographic adaptation in the Amazonian hyperdominant tree *Eperua falcata* Aubl. (Fabaceae)": Table S1 to S5

**Supplementary tables**

**Table S1.** Data description and Assembly statistics.

| | |
|---|---|
| Total number of paired reads | 2,332,116,952 reads |
| Number of paired reads in Lib 1 (Régina Terra-firme) | 567,889,784 |
| Number of paired reads in Lib 2 (Régina Bottomland) | 559,702,812 |
| Number of paired reads in Lib 3 (Laussat Terra-firme) | 494,014,536 |
| Number of paired reads in Lib 4 (Laussat Bottomland) | 710,509,820 |
| Total number of assembled contigs | 325,249 |
| Total assembly length | 249,308,901 bases |
| Mean depth | 57.81X |

**Table S2.** SNP detection. See also **Fig. S4**.

| | |
|---|---|
| Total number of bi-allelic SNPs | 97,062 |
| - Insertions-deletions | 2,954 |
| - Substitutions | 94,108 |
| o Transition | 60,483 (64% of substitutions) |
| o Transversions | 33,625 (36% of substitutions) |
| Ti/Tv | 1.80 |
| Mean SNP density (within polymorphic contigs) | 0.003/base (sd=0.004) |

**Table S3.** Summary of the distribution of normalised molecular diversity $\pi_L$ values in each sub-population. N = number of contigs carrying at least one polymorphism.

| Pop | N | 2.5% quantile | Median | 97.5% quantile |
|---|---|---|---|---|
| Régina terra firme | 17970 | 4.20E-05 | 3.09E-04 | 3.71E-03 |
| Régina bottomland | 18098 | 4.22E-05 | 3.09E-04 | 3.72E-03 |
| Laussat terra firme | 17280 | 4.49E-05 | 3.11E-04 | 3.45E-03 |
| Laussat bottomland | 19545 | 4.11E-05 | 3.16E-04 | 3.83E-03 |

**Table S4.** Genome-wide parameters estimated by the Bayesian model: $\mu r$ is the global mean of differentiation (over-all markers) at regional scale (i.e. between study sites), $\mu l_{Régina}$ and $\mu l_{Laussat}$ are the global mean of differentiation at local scale (i.e. between microhabitats within sites).

| | 2.5%CI | 25%CI | Median | 75%CI | 97.5%CI |
|---|---|---|---|---|---|
| $\mu r$ (subset1) | -3.69 | -3.55 | -3.5 | -3.41 | -3.2 |
| $\mu r$ (subset2) | -3.6 | -3.54 | -3.49 | -3.34 | -3.22 |
| $\mu r$ (subset3) | -4.04 | -3.88 | -3.55 | -3.28 | -3.14 |
| $\mu r$ (subset4) | -3.58 | -3.5 | -3.46 | -3.4 | -3.27 |
| $\mu r$ (subset5) | -3.9 | -3.75 | -3.58 | -3.44 | -3.36 |
| $\mu r$ (subset6) | -4 | -3.94 | -3.55 | -3.25 | -2.86 |
| $\mu r$ (subset7) | -3.72 | -3.64 | -3.45 | -3.38 | -3.33 |
| $\mu r$ (subset8) | -3.68 | -3.55 | -3.22 | -2.91 | -2.84 |
| $\mu r$ (subset9) | -3.66 | -3.58 | -3.49 | -3.36 | -3.25 |
| $\mu r$ (subset10) | -3.56 | -3.46 | -3.36 | -3.18 | -3.04 |
| $\mu l$ *Régina* (subset1) | -3.48 | -3.47 | -3.46 | -3.46 | -3.45 |
| $\mu l$ *Régina* (subset2) | -3.5 | -3.49 | -3.48 | -3.47 | -3.46 |
| $\mu l$ *Régina* (subset3) | -3.51 | -3.5 | -3.5 | -3.49 | -3.48 |
| $\mu l$ *Régina* (subset4) | -3.54 | -3.53 | -3.52 | -3.52 | -3.51 |
| $\mu l$ *Régina* (subset5) | -3.52 | -3.5 | -3.5 | -3.49 | -3.48 |
| $\mu l$ *Régina* (subset6) | -3.54 | -3.53 | -3.52 | -3.52 | -3.5 |
| $\mu l$ *Régina* (subset7) | -3.49 | -3.48 | -3.47 | -3.46 | -3.45 |
| $\mu l$ *Régina* (subset8) | -3.51 | -3.5 | -3.49 | -3.49 | -3.48 |
| $\mu l$ *Régina* (subset9) | -3.52 | -3.51 | -3.5 | -3.5 | -3.49 |
| $\mu l$ *Régina* (subset10) | -3.52 | -3.51 | -3.5 | -3.49 | -3.48 |
| $\mu l$ *Laussat* (subset1) | -3.48 | -3.47 | -3.46 | -3.45 | -3.44 |
| $\mu l$ *Laussat* (subset2) | -3.49 | -3.48 | -3.47 | -3.46 | -3.45 |
| $\mu l$ *Laussat* (subset3) | -3.5 | -3.49 | -3.48 | -3.48 | -3.46 |
| $\mu l$ *Laussat* (subset4) | -3.52 | -3.51 | -3.5 | -3.5 | -3.49 |
| $\mu l$ *Laussat* (subset5) | -3.5 | -3.49 | -3.48 | -3.48 | -3.46 |
| $\mu l$ *Laussat* (subset6) | -3.52 | -3.51 | -3.51 | -3.5 | -3.49 |
| $\mu l$ *Laussat* (subset7) | -3.48 | -3.47 | -3.47 | -3.46 | -3.45 |
| $\mu l$ *Laussat* (subset8) | -3.49 | -3.48 | -3.47 | -3.47 | -3.45 |
| $\mu l$ *Laussat* (subset9) | -3.5 | -3.49 | -3.48 | -3.48 | -3.47 |
| $\mu l$ *Laussat* (subset10) | -3.51 | -3.5 | -3.49 | -3.48 | -3.47 |

**Table S5.** Enrichment tests at microgeographic scale. The table lists the GO-terms significantly enriched in the subset of predicted genes neighboring outlier SNPs (<500kb) compared to all predicted genes ($X^2$ pvalue < 5%).

| GO_ID | GO_name | pvalue | Prop. in the entire dataset (%) | Prop. in the subset of genes neighboring outlier SNPs (%) |
|---|---|---|---|---|
| GO:0008409 | 5'-3' exonuclease activity | 0.001 | 0.015 | 0.943 |
| GO:0004534 | 5'-3' exoribonuclease activity | 0 | 0.003 | 0.943 |
| GO:0047617 | acyl-CoA hydrolase activity | 0 | 0.012 | 0.943 |
| GO:0006637 | acyl-CoA metabolic process | 0.001 | 0.018 | 0.943 |
| GO:0004013 | adenosylhomocysteinase activity | 0 | 0.012 | 0.943 |
| GO:0052646 | alditol phosphate metabolic process | 0.009 | 0.027 | 0.943 |
| GO:0042886 | amide transport | 0.005 | 0.823 | 3.774 |
| GO:0019202 | amino acid kinase activity | 0.001 | 0.018 | 0.943 |
| GO:0019439 | aromatic compound catabolic process | 0.006 | 0.201 | 1.887 |
| GO:1901136 | carbohydrate derivative catabolic process | 0 | 0.088 | 2.83 |
| GO:0009056 | catabolic process | 0.005 | 1.22 | 4.717 |
| GO:0003824 | catalytic activity | 0.025 | 24.176 | 33.962 |
| GO:0051641 | cellular localization | 0.008 | 0.872 | 3.774 |
| GO:0070727 | cellular macromolecule localization | 0.001 | 0.665 | 3.774 |
| GO:0042219 | cellular modified amino acid catabolic process | 0.003 | 0.021 | 0.943 |
| GO:0044270 | cellular nitrogen compound catabolic process | 0.001 | 0.149 | 1.887 |
| GO:0034613 | cellular protein localization | 0.001 | 0.665 | 3.774 |
| GO:0030131 | clathrin adaptor complex | 0.04 | 0.043 | 0.943 |
| GO:0016289 | CoA hydrolase activity | 0.013 | 0.03 | 0.943 |
| GO:0048475 | coated membrane | 0.007 | 0.204 | 1.887 |
| GO:0048037 | cofactor binding | 0.049 | 1.205 | 3.774 |
| GO:0008234 | cysteine-type peptidase activity | 0.045 | 0.314 | 1.887 |
| GO:0000910 | cytokinesis | 0.013 | 0.03 | 0.943 |
| GO:0000290 | deadenylation-dependent decapping of nuclear-transcribed mRNA | 0.001 | 0.015 | 0.943 |
| GO:0003006 | developmental process involved in reproduction | 0.032 | 0.04 | 0.943 |
| GO:0004143 | diacylglycerol kinase activity | 0.003 | 0.021 | 0.943 |
| GO:0051649 | establishment of localization in cell | 0.003 | 0.762 | 3.774 |
| GO:0045184 | establishment of protein localization | 0.004 | 0.805 | 3.774 |

| | | | | |
|---|---|---|---|---|
| GO:0004532 | exoribonuclease activity | 0.04 | 0.043 | 0.943 |
| GO:0016896 | exoribonuclease activity, producing 5'-phosphomonoesters | 0.04 | 0.043 | 0.943 |
| GO:0005542 | folic acid binding | 0 | 0.009 | 0.943 |
| GO:0010154 | fruit development | 0 | 0.009 | 0.943 |
| GO:0004349 | glutamate 5-kinase activity | 0 | 0.009 | 0.943 |
| GO:0046168 | glycerol-3-phosphate catabolic process | 0.001 | 0.018 | 0.943 |
| GO:0004367 | glycerol-3-phosphate dehydrogenase [NAD+] activity | 0.001 | 0.018 | 0.943 |
| GO:0009331 | glycerol-3-phosphate dehydrogenase complex | 0.003 | 0.021 | 0.943 |
| GO:0006072 | glycerol-3-phosphate metabolic process | 0.009 | 0.027 | 0.943 |
| GO:1901658 | glycosyl compound catabolic process | 0.001 | 0.015 | 0.943 |
| GO:0046700 | heterocycle catabolic process | 0.001 | 0.149 | 1.887 |
| GO:0016788 | hydrolase activity, acting on ester bonds | 0.028 | 1.574 | 4.717 |
| GO:0016801 | hydrolase activity, acting on ether bonds | 0 | 0.012 | 0.943 |
| GO:0006886 | intracellular protein transport | 0 | 0.601 | 3.774 |
| GO:0046907 | intracellular transport | 0.003 | 0.756 | 3.774 |
| GO:0009057 | macromolecule catabolic process | 0.025 | 0.628 | 2.83 |
| GO:0033036 | macromolecule localization | 0.021 | 1.031 | 3.774 |
| GO:0030117 | membrane coat | 0.007 | 0.204 | 1.887 |
| GO:0072341 | modified amino acid binding | 0.001 | 0.018 | 0.943 |
| GO:0051287 | NAD binding | 0.012 | 0.229 | 1.887 |
| GO:0043628 | ncRNA 3'-end processing | 0 | 0.003 | 0.943 |
| GO:0071705 | nitrogen compound transport | 0.023 | 1.046 | 3.774 |
| GO:0034655 | nucleobase-containing compound catabolic process | 0 | 0.122 | 1.887 |
| GO:0009164 | nucleoside catabolic process | 0.001 | 0.015 | 0.943 |
| GO:1901265 | nucleoside phosphate binding | 0.029 | 9.375 | 16.038 |
| GO:0000166 | nucleotide binding | 0.029 | 9.375 | 16.038 |
| GO:0006730 | one-carbon metabolic process | 0.001 | 0.015 | 0.943 |
| GO:1901361 | organic cyclic compound catabolic process | 0.006 | 0.201 | 1.887 |
| GO:1901575 | organic substance catabolic process | 0.002 | 1.083 | 4.717 |
| GO:1901565 | organonitrogen compound catabolic process | 0.016 | 0.576 | 2.83 |
| GO:0046434 | organophosphate catabolic process | 0.009 | 0.027 | 0.943 |
| GO:0016903 | oxidoreductase activity, acting on the aldehyde or oxo group of donors | 0.005 | 0.195 | 1.887 |
| GO:0016624 | oxidoreductase activity, acting on the aldehyde or oxo group of donors, disulfide as acceptor | 0.04 | 0.043 | 0.943 |
| GO:0004591 | oxoglutarate dehydrogenase (succinyl-transferring) activity | 0.001 | 0.018 | 0.943 |
| GO:0015833 | peptide transport | 0.005 | 0.817 | 3.774 |
| GO:0016774 | phosphotransferase activity, carboxyl group as acceptor | 0.009 | 0.027 | 0.943 |

| | | | | |
|---|---|---|---|---|
| GO:0009791 | post-embryonic development | 0.032 | 0.04 | 0.943 |
| GO:0097354 | prenylation | 0.001 | 0.018 | 0.943 |
| GO:0004659 | prenyltransferase activity | 0.025 | 0.037 | 0.943 |
| GO:0006561 | proline biosynthetic process | 0 | 0.012 | 0.943 |
| GO:0006560 | proline metabolic process | 0.001 | 0.015 | 0.943 |
| GO:0018344 | protein geranylgeranylation | 0 | 0.009 | 0.943 |
| GO:0004661 | protein geranylgeranyltransferase activity | 0 | 0.006 | 0.943 |
| GO:0007205 | protein kinase C-activating G-protein coupled receptor signaling pathway | 0.003 | 0.021 | 0.943 |
| GO:0008104 | protein localization | 0.006 | 0.848 | 3.774 |
| GO:0018342 | protein prenylation | 0.001 | 0.018 | 0.943 |
| GO:0008318 | protein prenyltransferase activity | 0 | 0.012 | 0.943 |
| GO:0015031 | protein transport | 0.004 | 0.784 | 3.774 |
| GO:0006152 | purine nucleoside catabolic process | 0 | 0.012 | 0.943 |
| GO:0046130 | purine ribonucleoside catabolic process | 0 | 0.012 | 0.943 |
| GO:0072523 | purine-containing compound catabolic process | 0 | 0.012 | 0.943 |
| GO:0004663 | Rab geranylgeranyltransferase activity | 0 | 0.006 | 0.943 |
| GO:0048608 | reproductive structure development | 0.032 | 0.04 | 0.943 |
| GO:0061458 | reproductive system development | 0.032 | 0.04 | 0.943 |
| GO:0042454 | ribonucleoside catabolic process | 0.001 | 0.015 | 0.943 |
| GO:0090503 | RNA phosphodiester bond hydrolysis, exonucleolytic | 0.04 | 0.043 | 0.943 |
| GO:0019510 | S-adenosylhomocysteine catabolic process | 0 | 0.012 | 0.943 |
| GO:0046498 | S-adenosylhomocysteine metabolic process | 0 | 0.012 | 0.943 |
| GO:0048316 | seed development | 0 | 0.009 | 0.943 |
| GO:0003697 | single-stranded DNA binding | 0.048 | 0.046 | 0.943 |
| GO:0036094 | small molecule binding | 0.003 | 9.851 | 18.868 |
| GO:0034472 | snRNA 3'-end processing | 0 | 0.003 | 0.943 |
| GO:0016073 | snRNA metabolic process | 0 | 0.009 | 0.943 |
| GO:0016180 | snRNA processing | 0 | 0.009 | 0.943 |
| GO:0044273 | sulfur compound catabolic process | 0.003 | 0.021 | 0.943 |
| GO:0006790 | sulfur compound metabolic process | 0.025 | 0.271 | 1.887 |
| GO:0030976 | thiamine pyrophosphate binding | 0.04 | 0.043 | 0.943 |
| GO:0035383 | thioester metabolic process | 0.001 | 0.018 | 0.943 |
| GO:0016790 | thiolester hydrolase activity | 0 | 0.052 | 1.887 |
| GO:0016802 | trialkylsulfonium hydrolase activity | 0 | 0.012 | 0.943 |
| GO:0034477 | U6 snRNA 3'-end processing | 0 | 0.003 | 0.943 |
| GO:0019842 | vitamin binding | 0.001 | 0.357 | 2.83 |
