## Supplementary material for "Genomic and phenotypic divergence unveil microgeographic adaptation in the Amazonian hyperdominant tree *Eperua falcata* Aubl. (Fabaceae)": File S1

### Background biological and environmental description of populations and sites

A subset of the trees sampled for genomic analyses was also described for growth and functional traits; soil properties were also described at the foot of each tree in that subsample, to analyse the environmental determinants of those trait values. These data are under analysis and will be the subject of a separate paper (Scotti et al., in prep.).

We present here an overview of population and site properties as a description of the samples we have analysed.

#### 1. Soil properties

Habitats differ mainly for water table depth, as indicated by the following plot. Here, water table classes, based on the depth of the water table at time of soil sample collection, are used (0 = no water table found; 1 = water table at 0.6-1.2 m below ground; 2 = water table at 0.1-0.6 m below ground; 3 = water table above 0.1 m):

Laussat has a typical bottomland – *terra firme* profile, with flooding in the bottomland and a deep water table on the hilltop; at Régina, consistently with the more rugged landscape, there is no true flooded area, and water table depth varies widely in the bottomland, without reaching truly flooded classes, with rare exceptions.

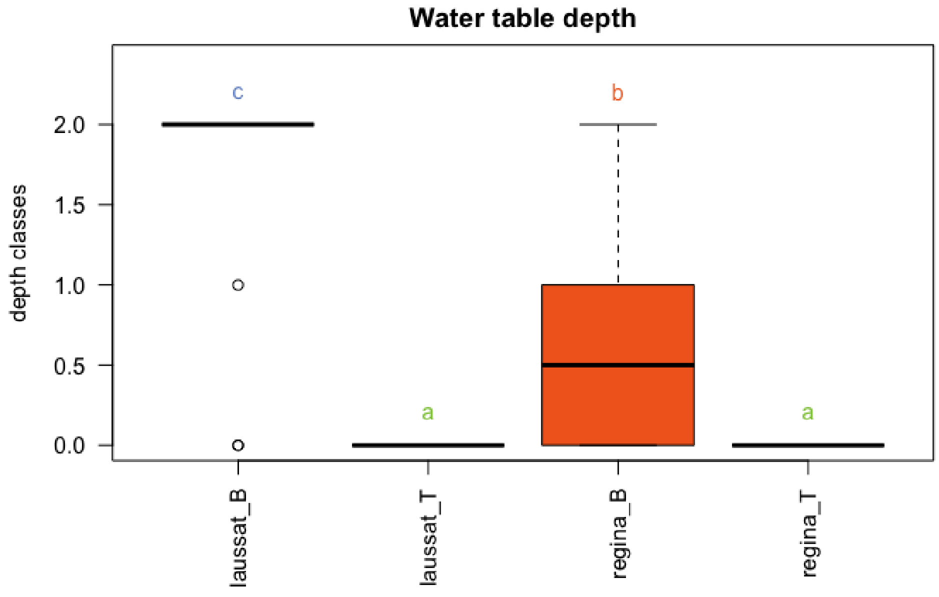

General soil composition differs between sites and habitats, as shown by the following plot, displaying the proportion of trees on each soil (red = ferrallitic soil, dark grey = hydromorphic soil, wheat = podzol soil, shades of brown = intermediate podzol – ferrallitic soil):

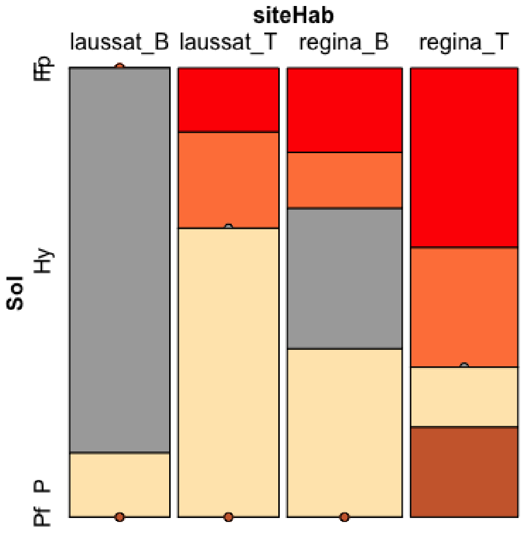


The Laussat pair displays a stark contrast between hydromorphic soil in the seasonally flooded habitat and podzol on the higher ground; the situation is less clear at Régina, where both habitats have more ferrallitic and less hydromorphic soil, with large differences mostly in the intermediate ferrallitic - podzol classes.

Soils across the two sites are similar for C:N ratio, cations and pH; the tow sites differ in sand and silt composition, and differences between habitats within site appear for phosphorus content and, partially, sand content.

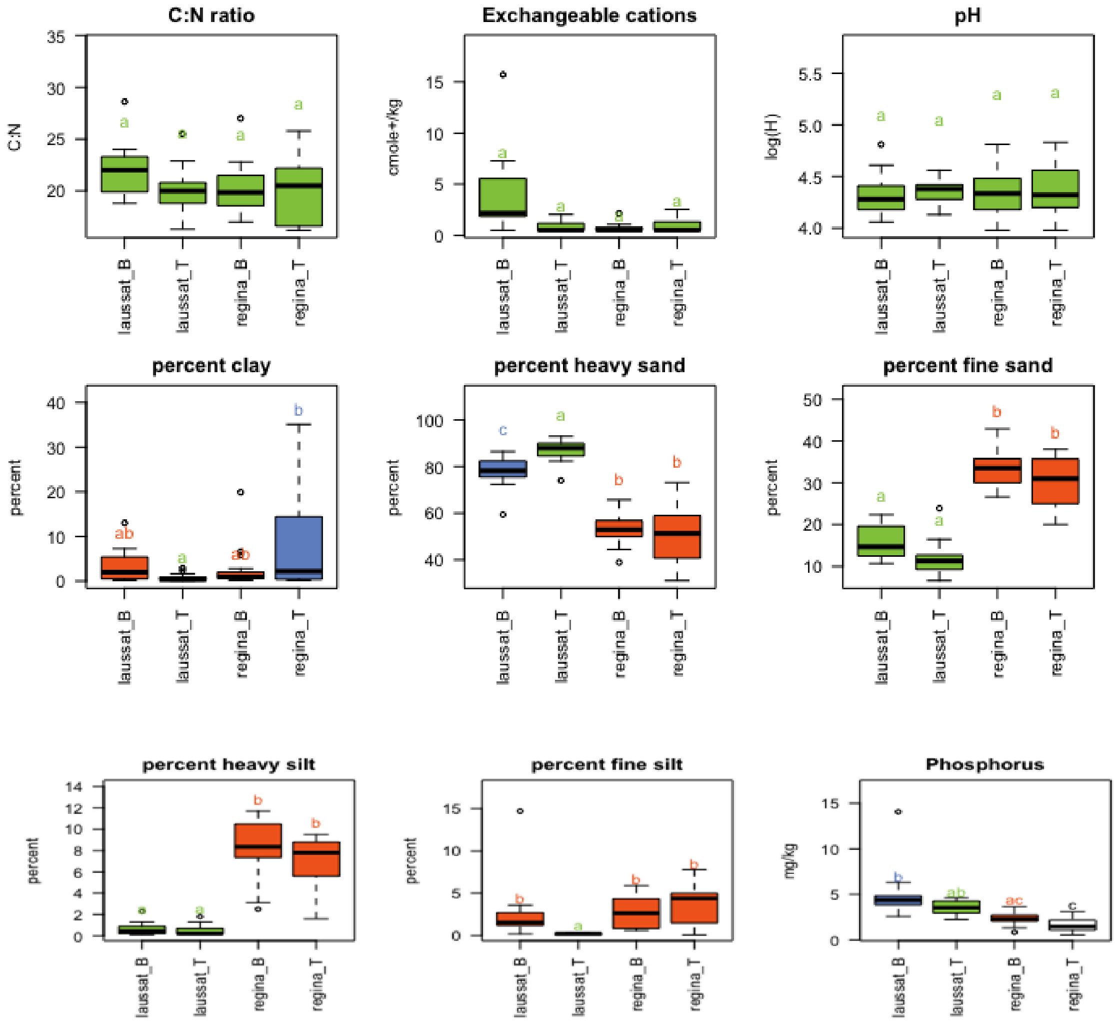

#### 2. Tree properties

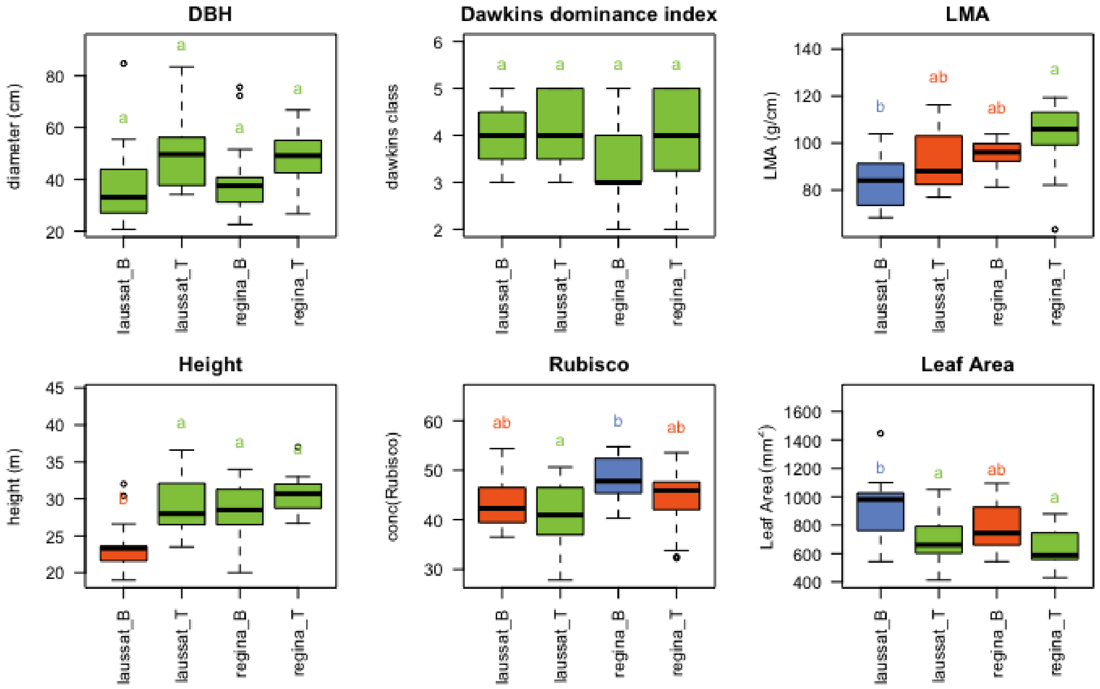
