## Supplementary material for "Genomic and phenotypic divergence unveil microgeographic adaptation in the Amazonian hyperdominant tree *Eperua falcata* Aubl. (Fabaceae)": File S2

### Bioinformatics pipeline

#### Read pre-processing

Paired-end reads (100 nt) were pre-processed (cleaned and filtered) before assembly. Individual bases of low quality were masked using the 'Fastq_masker' tool of the suite fastx_toolkit (http://hannonlab.cshl.edu/fastx_toolkit) under a quality threshold of 25. Reads-ends were trimmed using Fastq_trimming' (fastx_toolkit) by trimming the first and last ten bases of each reads (resulting in reads of 80 nt). Reads were further sorted and orphan reads were excluded using Sickle (Joshi & Fas, 2011) with the following parameters: minimum per base quality threshold = 25 and minimum read length = 70. The quality of the cleaned reads was assessed using 'fastqc'. Reads pre-processing resulted in 2,332,116,952 high-quality reads (**Table S1**). Cleaned reads have been deposited on the European Nucleotide Archive (ENA, EMBL), under project accession code PRJEB9879, sample accession codes ERS791957, ERS791958, ERS791959, ERS791960.

#### *De novo* assembly and mapping

*E. falcata* is a non-model species with no reference genome. Its genome size is unknown, but published data from twenty-two species (compiled at https://cvalues.science.kew.org/) suggest that genome sizes in the Caesalpiniaceae sub-family are comprised between 0.6 and 2 Gbp (mean size = 0.90 Gbp) (Bennett, Leitch, & Hanson, 1998; Ohri, Bhargava, & Chatterjee, 2004; Ohri, Kumar, & Pal, 1986; Ohri & Kumar, 1986; Sliwinska, Pisarczyk, Pawlik, & Galbraith, 2009). Sequence data acquired prior to this study revealed a diploid status (Audigeos, Brousseau, Traissac, Scotti-Saintagne, & Scotti, 2013).

Cleaned reads were assembled into a crude *de novo* genome assembly which constitutes a large new pan-genomic resource. Because classical assemblers do not easily support the numerous polymorphisms contained in libraries of pooled individuals (resulting in the segregation of reads from different populations pools into different assembled contigs and scaffolds), the sequenced libraries were *de novo* assembled using RayMeta (Boisvert, Raymond, Godzaridis, Laviolette, & Corbeil, 2012), an assembler originally developed for meta-communities. The different libraries were assembled together in order to maximize the coverage of the *de novo* assembly, with a default k-mer size of 21 which optimized the memory used and the computation time (the minimum k value allowed by Ray being 15). Even if large value of k could be preferred, long k-mer increase the chances of error and require much more memory (Chikhi & Medvedev, 2014). The assembled contigs were then aligned together in order to re-align redundant contigs using CD-HIT-EST (Fu, Niu, Zhu, Wu, & Li, 2012; W. Li & Godzik, 2006) with a sequence identity threshold of 95%. Non-redundant contigs were retained and used as reference to map *a posteriori* each library separately using bwa (H. Li & Durbin, 2009): 'bwa aln' and 'bwa sampe' (only paired reads were mapped). The .sam files (one per library) were converted into .bam, sorted, indexed and merged into a unique .mpileup file using the samtools suite (H. Li et al., 2009): 'samtools view', 'samtools sort', 'samtools index' and 'samtools mpileup'.

**Structural and functional annotation**

The crude pangenomic reference was annotated with the suite implemented in Maker mpi v. 2.31.8 (Cantarel et al., 2008) and associated tools. In summary, repetitive elements were detected by repeatMasker v.4.0.3 (http://www.repeatmasker.org/) and *ab initio* gene prediction was realized by Augustus v.2.6.1 (Stanke et al., 2006) at the scaffold level using both the reference transcriptome of *E. falcata* (Brousseau et al., 2014) and a composite proteome reference composed of all published reference proteomes of Fabaceae (*Cajanus cajan*, *Cicer arietinum*, *Glycine max*, *Lupinus angustifolius*, *Medicago truncatula*, *Phaseolus angularis*, *Phaseolus vulgaris* and *Vigna radiata*) plus the reference proteomes of *Arabidopsis thaliana*. The predicted proteins were further BLASTed against the non-redundant protein database with NCBI BLASTp v.2.6.0 retaining the first best hit only. At last, gene ontology terms of each predicted gene were retrieved using Interproscan 5.21.60 (Jones et al., 2014).

**SNP detection and Filtering**

The absolute abundance of different nucleotides was counted for each assembled position using the PoPoolation2 mpileup2sync.jar script (Kofler, Pandey, & Schlötterer, 2011). The .java.sync output file containing nucleotide counts for each position of the assembly was post-processed through custom perl scripts.

(1) Monomorphic sites were discarded and rare SNPs rare were masked (replaced by 'N') under a minimum allele frequency (maf) threshold of 0.05, all libraries confounded (meaning that a minimum number of 10 alternative variants out of 200 sequenced gametes in the whole dataset was required for a polymorphism to be retained.

(2) Among the remaining SNPs, only bi-allelic polymorphisms with a total maximum coverage of 200X over all libraries to avoid SNPs caused by potential misassembly of paralogs into the same contigs and scaffolds.

Given the minimum per-base phred-score threshold of 25 used, the corresponding probability of incorrect base call would be $P = 10^{(-25/10)} \approx 0.32\%$. Consequently, the probability of calling X times a wrong alternative variant (*i.e.*, of detecting a false SNP) with a minimum depth of 200X and a maf of 0.05 would be below $10^{-25}$.

(3) Bi-allelic SNPs with library-level coverage < 20X were discarded, resulting in a SNP dataset composed of SNPs with coverage > 20X within each of the four libraries.

(4) Contiguous SNPs were further filtered out to avoid polymorphisms caused by copy number variation (CNV). (4 and 5) SNPs located within repeat elements (detected by repeatMasker) as well as SNPs located in contigs with SNP density > 0.02 were filtered out, once again to avoid SNPs located in paralogs (coding sequence data in Audigeos et al. (2013) suggest that SNP density in *E. falcata* is on average around 0.03, so our threshold for genomic data (0.02) seems to be reasonable, although conservative). Linkage Disequilibrium has been shown to decay very quickly with distance (*i.e.*, it falls below $r^2 = 0.2$ within 100-200 bp) in *E. falcata* coding sequences) (Audigeos et al., 2013).
