## Supplementary material for "Genomic and phenotypic divergence unveil microgeographic adaptation in the Amazonian hyperdominant tree *Eperua falcata* Aubl. (Fabaceae)": File S3

### Bayesian Model specifications

The Bayesian model was encoded in stan (Carpenter et al., 2016) and compiled into C++ using Rstan. Because Bayesian inference is computationally expensive, the model was applied to subsets of 10,000 SNPs on a Dell Power Edge CG220 Intel Xeon (CPU E5-2670 v2 @2.50Ghz) cluster node with parallelization on 5 CPU (with parameters: virtual memory vmem = 25G, virtual memory limit h_vmem=30G and parallel_fill = 5) and explicit chains parallelization in R with the package 'parallel' using the function 'stan' : 1000 iterations, 2 chains. The warmup period was adapted by Stan with the option 'step-size adaptation' (which it is by default). Chain convergence was assessed with the function 'traceplot'. Draws were extracted using the function 'extract' including the statement 'permuted=TRUE' to only save the draws after the warmup period with permutation so that they can be analyzed directly without concerns of autocorrelation. Further details on the functions 'stan', 'extract' and 'traceplot' can be found at https://mc-stan.org/rstan/reference/stan.html, https://mc-stan.org/rstan/reference/stanfit-method-extract.html and https://mc-stan.org/rstan/reference/stanfit-method-traceplot.html, respectively. Both dataset hashing and explicit parallelization optimized computation times and the model ran in ~ 4 hours / subset. Outputs were post-processed with R, including chain convergence checking and eBP values inference from the posterior distribution of locus-specific parameters. Due to the large number of SNPs screened, a correction for multiple-testing was applied on eBP values with the package 'fdrtool' with an arbitrary FDR threshold of 10%. The proportions of outliers under selection was compared between geographic scales with chi-squared tests ('prop.test' function). Enrichment tests were further conducted through $X^2$ tests in order to test (i) the enrichment of outliers located within exons, within a predicted gene and in the neighborhood (< 5 kb) of a predicted gene compared to the whole SNP dataset, as well as (ii) the enrichment of GO terms between predicted genes neighboring outlier SNPs compared to all predicted genes of the entire reference.
