## Supplementary material for "Genomic and phenotypic divergence unveil microgeographic adaptation in the Amazonian hyperdominant tree *Eperua falcata* Aubl. (Fabaceae)": File S4

### BayPass methodology & results

#### Overview

BayPass v.2.1. (Gautier, 2015) was applied to our pool-seq dataset using counts data as a complementary approach to test whether the analyzed SNPs were submitted to divergent adaptation across populations. This method first estimates the neutral genetic structuring though a genome-wide variance-covariance matrix ($\Omega$ whose precision matrix $\Omega^{-1}=\Delta$) and standardize allele frequencies ($\alpha^*_{i,j}$) for each marker $i$ within each population $j$ accordingly to finally estimate the extent of locus-specific genetic differentiation for each marker $i$ ($XtX_i$, Günther & Coop, 2013).

$XtX_i$ estimates thus consist in the overall genetic differentiation for each SNP between actual populations and the ancestral population corrected by the neutral genetic structuring across populations. This statistics allows identifying SNPs that are over-differentiated across all studied populations, but it does not allow distinguishing SNPs that are structured by the two binary and independent factors tested here -study sites and micro-habitats, nor to test for each factor independently.

#### Procedure

BayPass core model was first run on the real SNPs dataset with 15 pilot runs, a burning of 2500 followed by 5000 iterations and a thinning of 15.

```
g_baypass -outprefix wapa -nthreads 8 -npop 4 -gfile
/scratch/scratch_wapa/wapa_data.counts -poolsizefile wapa_data.pool -thin 15
-burnin 2500 -npilot 15 -pilotlength 5000
```

where the 'gfile' contains allele counts for each bi-allelic SNP within each population and the 'poolsizefile' describes the size of haploid sequences within each population pool, that is 40 for each pool.

The variance-covariance matrix (described in the file 'wapa_mat_omega.out') and the mean priors of the Beta distribution of the ancestral reference allele frequency ($a_\pi$ and $b_\pi$ described in the file 'wapa_summary_beta_params.out') estimated were further used to simulate a pseudo-observed dataset of 100000 SNPs (PODs) using the R function 'simulate.baypass':

```
simu=simulate.baypass(omega.mat=omega,nsnp=100000,sample.size=raw.data$NN,beta.pi=pi.beta.coef,pi.maf=0.05,suffix="wapa_pods")
```

BayPass was run a second time on PODs under the same parameters as for the real dataset:

```
g_baypass -outprefix wapa_pods -nthreads 8 -npop 4 -gfile /scratch/scratch_wapa/G.wapa_pods -thin 15 -burnin 2500 -npilot 15 -pilotlength 5000
```

Outputs were post-processed using R according to the user manual instructions. A sanity check was conducted by estimating the Forstner and Moonen distance (Förstner & Moonen, 2003) between omega covariance matrices estimated on real data and on PODs (function 'fmd.dist'). An arbitrary *XtXi* significance threshold of 99% was estimated using PODs to identify SNPs over-differentiated across all populations without consideration of the spatial scale considered (regional or microgeographic adaptive divergence). Two-sample Wilcoxon tests (function 'wilcox.test') were applied to compare *XtXi* between sets of SNPs identified as neutral and SNPs identified as outliers under divergent selection at each spatial scale (regional and microgeographic, respectively) through our custom Bayesian model.

#### Results

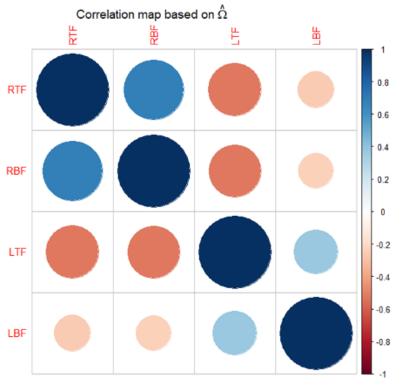


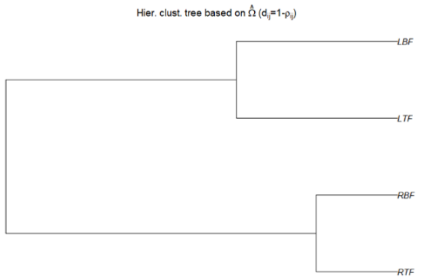


**<u>Variance-covariance matrix (Ω real).</u>** It confirmed the existence of a hierarchical neutral structuring across populations that is consistent to our expectations and $G_{ST}$ estimates between sites and micro-habitats within sites.

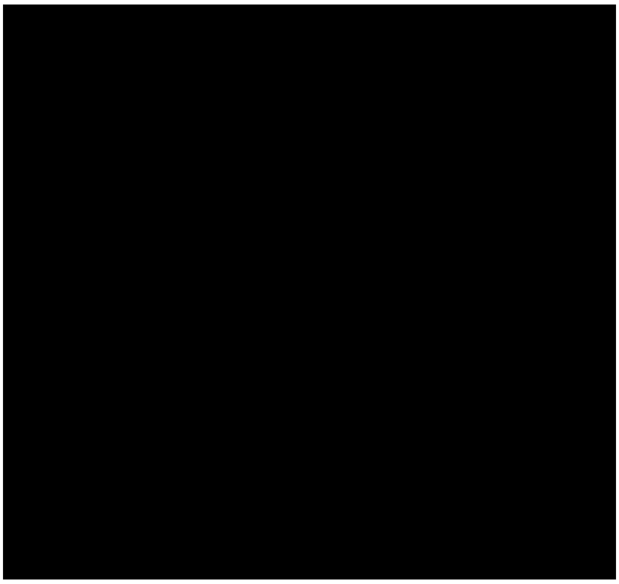

**<u>Sanity check</u>**. Comparing Ω between real data and PODs underpinned that pseudo-observed data are consistent with the hierarchical neutral structuring of the real dataset. Fmd distance (real , PODs)= 0.7245678.

(a)

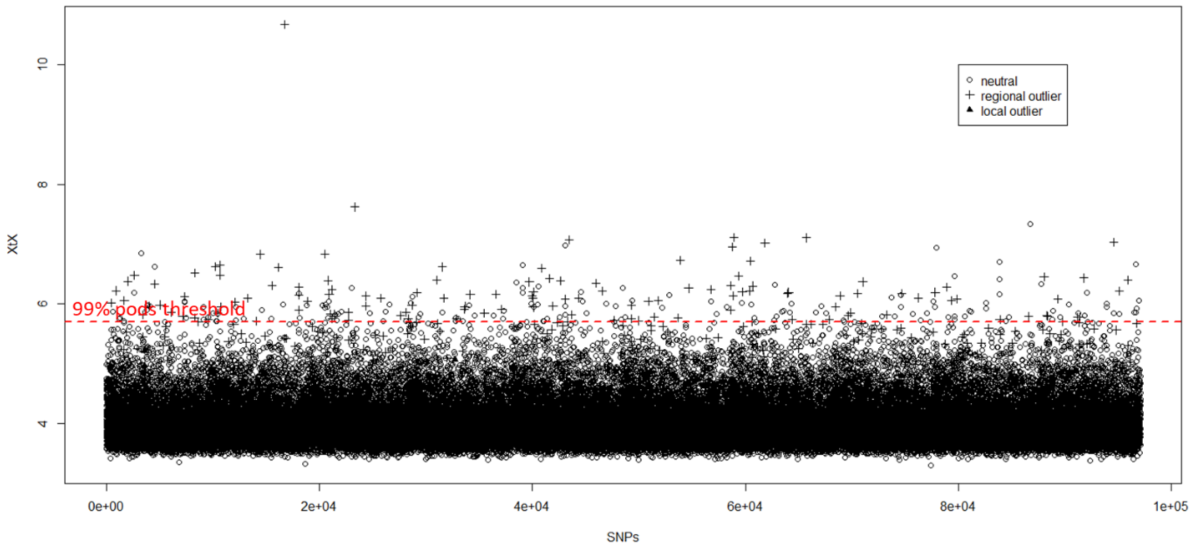


(b)

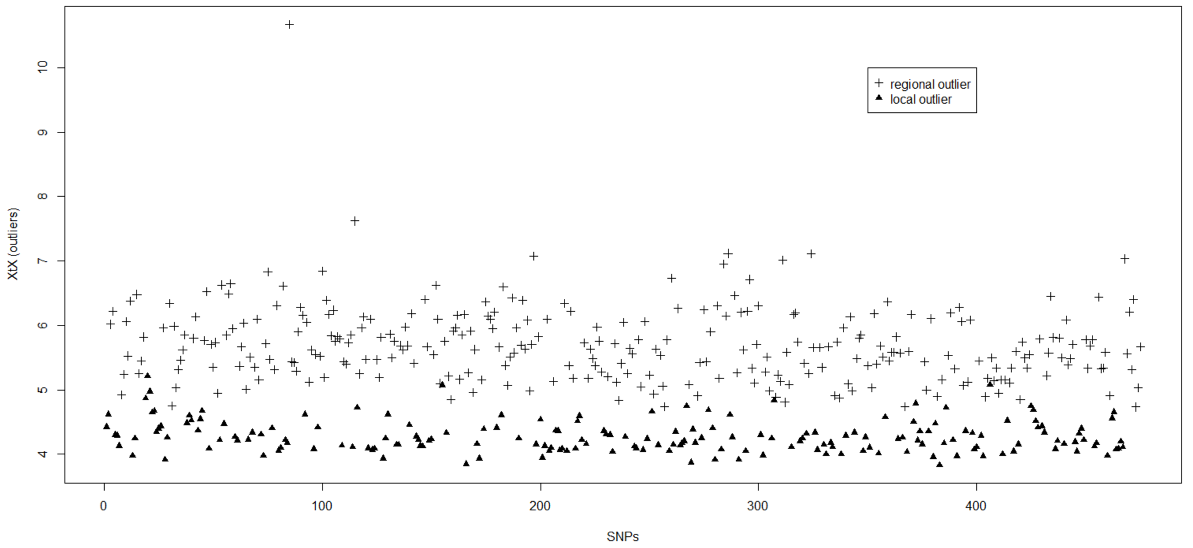


**Manhattan plots on $XtX_i$.** Estimates of SNP-specific genetic differentiation ($XtX_i$ statistics) for every SNPs (a) and for those detected as outliers by our custom Bayesian model only (b). Different symbols are used to distinguish neutral SNPs (○), outliers under divergent selection at regional scale (+) and outliers under divergent selection at microgeographic scale (▲).

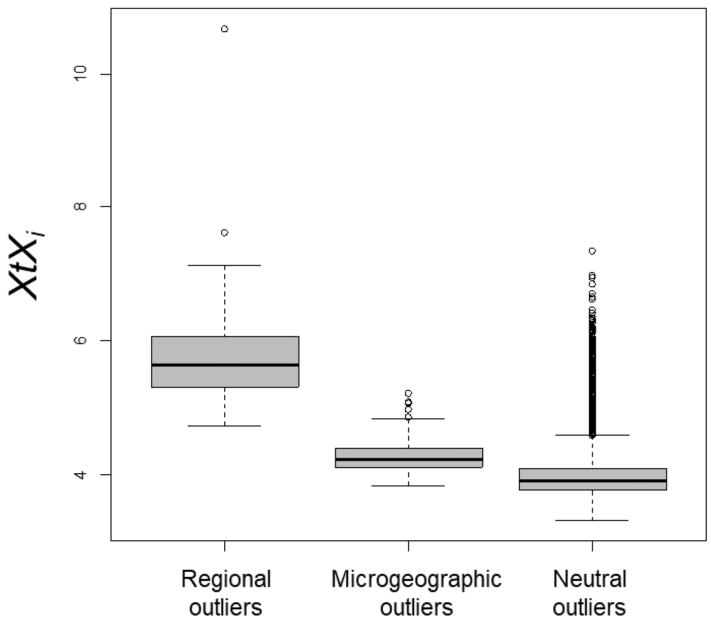


**<u>Comparison of *$XtX_i$* across sets of SNPs.</u>** The distribution of *$XtX_i$* was significantly lower for SNPs identified as neutral under our custom Bayesian model compared to SNPs identified as outliers at both regional and microgeographic scales. Results of Wilcoxon tests are provided below:

```
# Neutral SNPs vs outliers at regional scale
> wilcox.test(xtx.outliers.site,xtx.neutral,alternative="greater")
Wilcoxon rank sum test with continuity correction
data: xtx.outliers.site and xtx.neutral
W = 27866000, p-value < 2.2e-16
alternative hypothesis: true location shift is greater than 0

# Neutral SNPs vs outliers at microgeographic scale
> wilcox.test(xtx.outliers.micro,xtx.neutral,alternative="greater")
Wilcoxon rank sum test with continuity correction
data: xtx.outliers.micro and xtx.neutral
W = 14542000, p-value < 2.2e-16
alternative hypothesis: true location shift is greater than 0
```
