## Supplementary material for "Genomic and phenotypic divergence unveil microgeographic adaptation in the Amazonian hyperdominant tree *Eperua falcata* Aubl. (Fabaceae)": File S5

### G2D methodology & results

We applied the G2D test method developed by (Nielsen et al., 2009) to the identification of contigs displaying outstanding patterns of two- population allele frequencies at the SNPs they contain.

**Overview**

In brief, the method analyses the two-population (and therefore two-dimensional) Site Frequency Spectrum (2D-SFS), and is based on the estimation of the multinomial probability (defined by the *G2D* statistic) of observing a given combination of single-contig allele counts on a two-dimensional grid, based on the expected probabilities provided by the background genomic distribution of counts. Contigs with low likelihood are considered as departing from the neutral demographic genome-wide expectation. Because it is impossible to establish a theoretical distribution of G2D values for an arbitrary demographic model, the expected distribution of G2D values for a given population sample is obtained by simulation. First, a demographic model is fitted to the empirical data, and then a large number of data sets is simulated under the best model. The distribution of the G2D values obtained on the simulated data is used as a reference; the P-value associated to each empirical single-contig data is obtained by first computing the G2D statistic on the contig, and then comparing it to the simulation-based reference.

Note that the identity of ancestral and derived variants is unknown for *Eperua falcata*, and therefore we used the folded Site Frequency Spectrum ('fSDS') in all subsequent analyses.

**Demographic model**

We used a coalescent approach, combined with a maximum-likelihood analysis, as implemented in FastSimCoal version fsc251 (Excoffier, Dupanloup, Huerta-Sánchez, Sousa, & Foll, 2013; Excoffier & Foll, 2011), to identify the best demographic model for a 4D-fSFS. To keep the model's computational demands reasonable, a random sample of 3000 contigs was used in demographic modelling.

A simple coalescent demographic model was applied, with within-site populations merging first, and then regional populations. All populations (both current and ancestral) were allowed one free size change (expansion, contraction, or stability) before merging; gene flow was allowed both within sites and between sites. The maximum-likelihood algorithm implemented in FastSimCoal does not use priors, but ranges, of which only the lower boundary is meaningful (the upper boundary is only a reference, and the algorithm is allowed to increase the boundary at each cycle). Lower boundaries for all parameters were set at the minimum (i. e. 1 for population sizes and event times, 0 for migration rates and past-to-present population size ratios).

The best-likelihood model had the following parameter values:

| | | | | | |
|---|---|---|---|---|---|
| Current size, Régina Hilltop | 278 | Time of size change, Régina Hilltop | 14 | Size change ratio, Régina Hilltop | 2.94 |
| Current size, Régina Bottomland | 276 | Time of size change, Régina Bottomland | 129 | Size change ratio, Régina Bottomland | 0.16 |
| Current size, Laussat Hilltop | 489 | Time of size change, Laussat Hilltop | 7 | Size change ratio, Laussat Hilltop | 8.51 |
| Current size, Laussat Bottomland | 70 | Time of size change, Laussat Bottomland | 293 | Size change ratio, Laussat Bottomland | 0.62 |
| Merger time, Laussat populations | 460 | Time of size change, Laussat merged | 642 | Size change ratio, Laussat merged | 0.22 |
| Merger time, Régina populations | 340 | Time of size change, Régina merged | 623 | Size change ratio, Régina merged | 0.72 |
| Merger time, Régina and Laussat | 665 | Time of size change, Merged Régina and Laussat | 777 | Size change ratio, Merged Régina and Laussat | 3.99 |
| Migration rate, Régina-Laussat | 0.017 | | | | |
| Migration rate, Régina Hilltop-Bottmland | 0.357 | | | | |
| Migration rate, Laussat Hilltop-Bottomland | 0.204 | | | | |

**Density distribution of G2D-statistic for simulated and empirical 2D-fSFS tables**

These parameters were used to generate 100 simulations of 3000 sequences, to mimic the empirical data set. This provided 30000 sequences, that were used to construct 2D-fSFS tables for each of the two simulated sites, and to compute simulated values of the G2D-statistic for each contig.

To obtain G2D-statistic values from SFS tables, the custom G2DcalcMultiPop() R function (Appendix A) was used.

The same function was used for the computation of the G2D-statistic on empirical data.

**Identification of G2D outliers**

All the subsequent analyses and plotting activities were perfomed in R.

We search for within-site (that is, within Régina and within Laussat) G2D outliers (that is, contigs whose SNPs had an unlikely distribution in the 2D-fSFS, relative to the genomic expectation). To do this, we compared individual contig G2D values to the G2D distrubtion obtained from simulated data. Empirical G2D values higher than the 95%-th quantile of the distribution were deemed as significant at the α = 5% significance level.

The test returned six outlier contigs for Régina, and 4361 outliers for Laussat. This large difference in output may point to the demographic model not representing well the structure of the Laussat site (e.g., time of population size change is estimated at seven generations ago for Laussat Hilltop, and the estimation of current population size for Laussat Bottomland is only 70), as the distribution of empirical and simulated G2D values look very different for Laussat, but not for Régina:

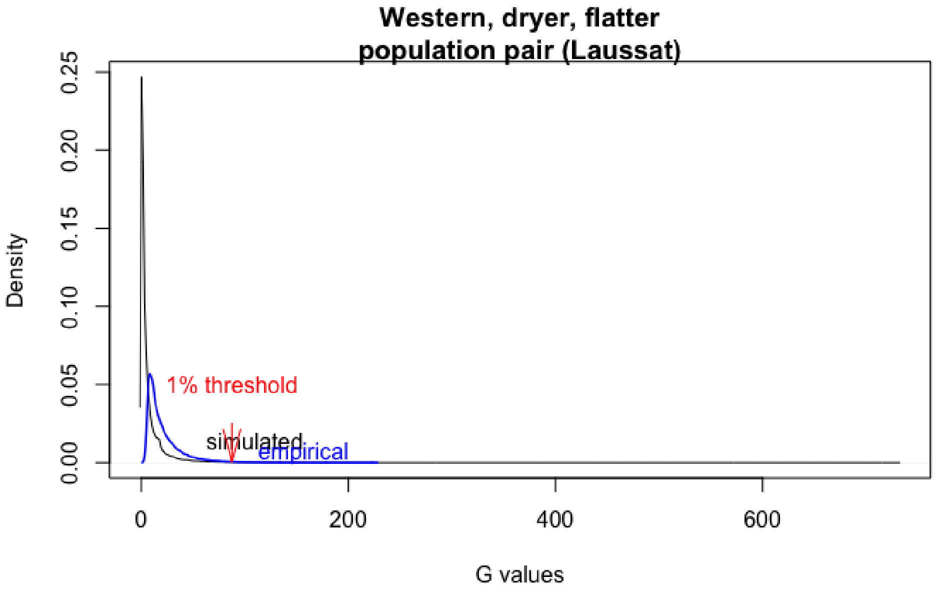


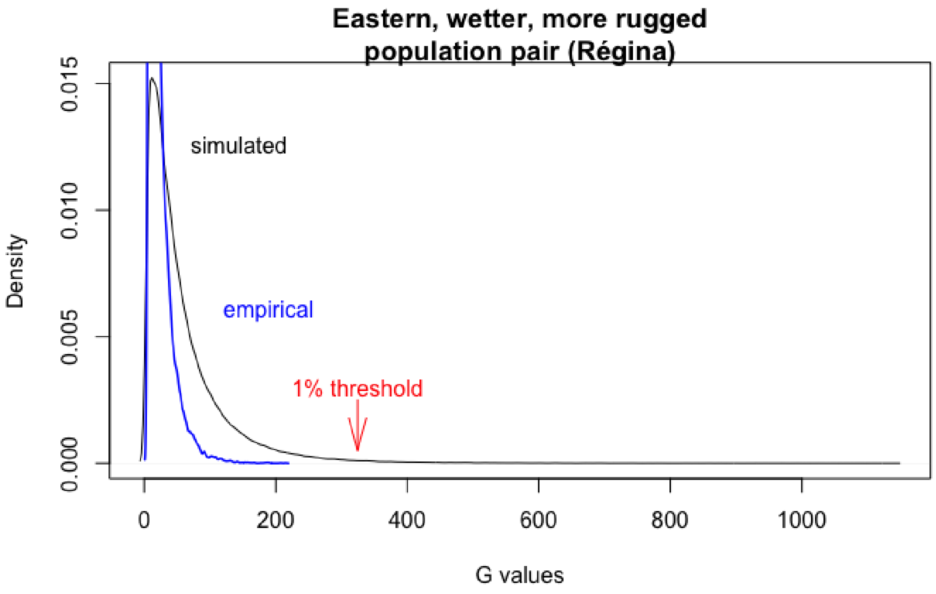


The six outlier contigs occurring at Régina also occurred at Laussat. The distribution of SNP 2D-fSFS for each of the six contigs, against the genome-wide backdrop, at the two sites is displayed below.

**contig_11657000001_Régina**

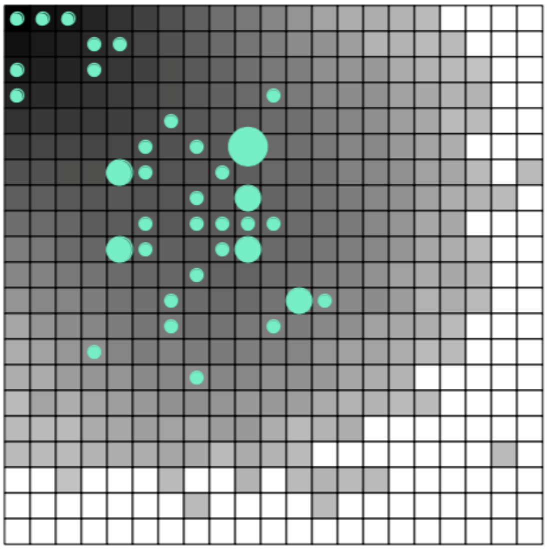

**contig_11657000001_Laussat**

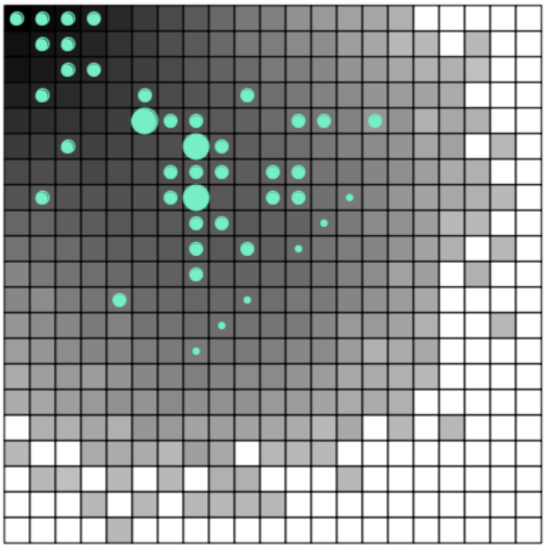

**contig_20678000007_Régina**

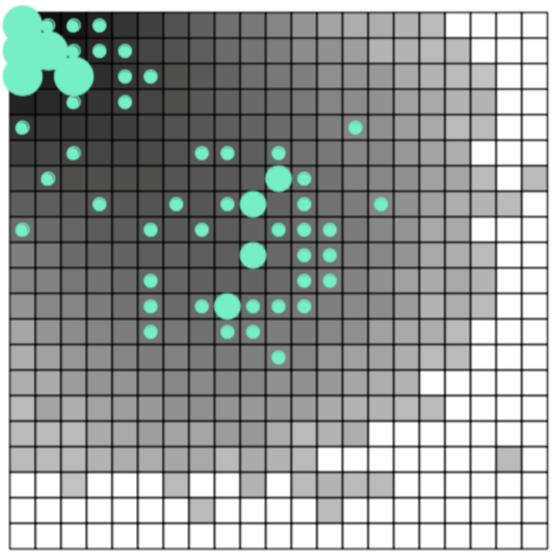

**contig_20678000007_Laussat**

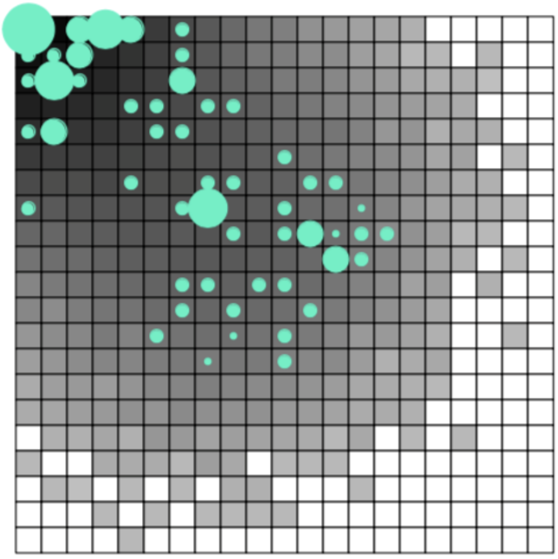

contig_27043000008_Régina

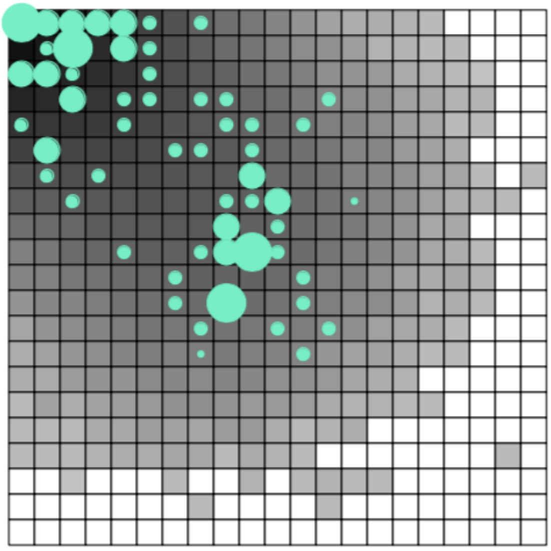

contig_27043000008_Laussat

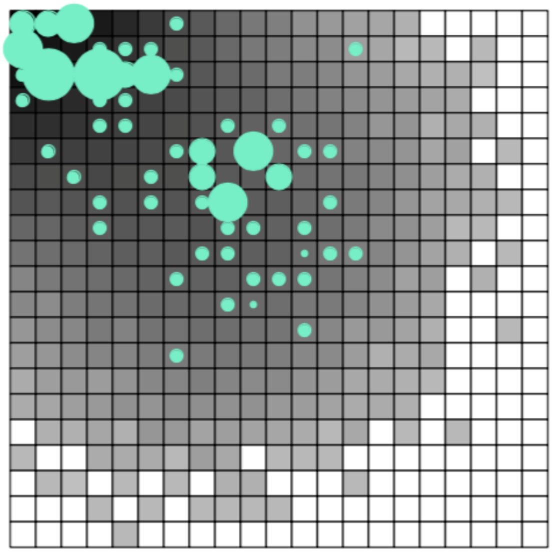

contig_29833000008_Régina

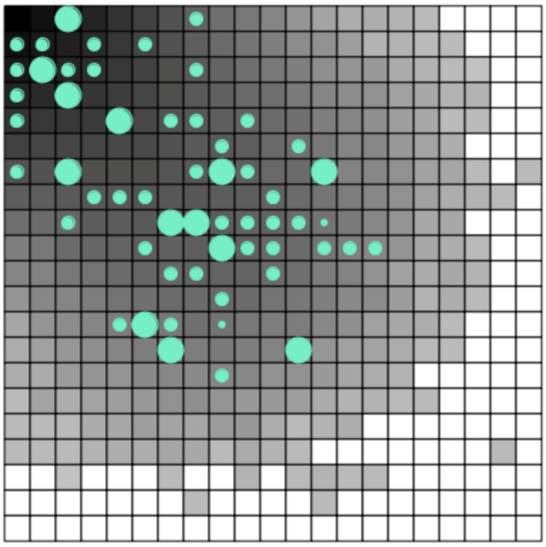

contig_29833000008_Laussat

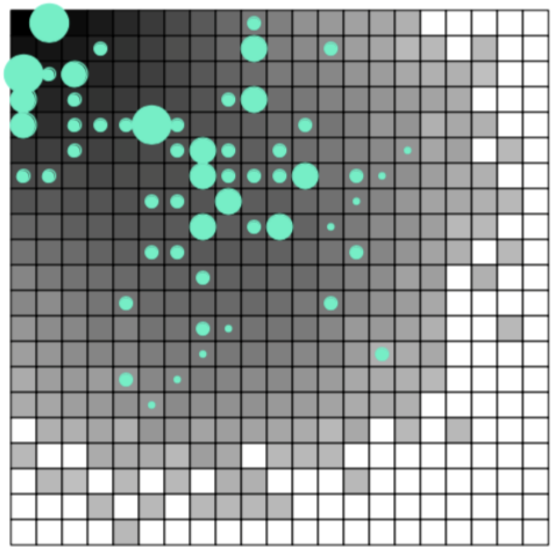

contig_5820000004_Régina

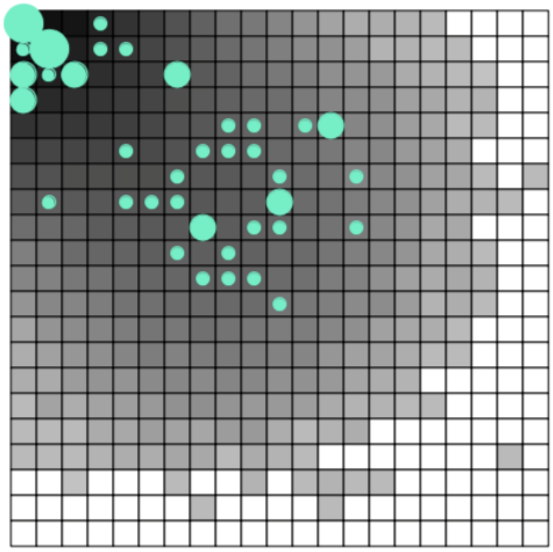

contig_5820000004_Laussat

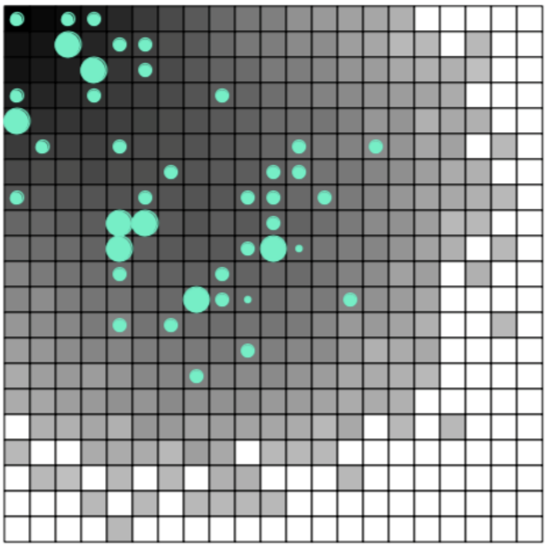

contig_9747000008_Régina

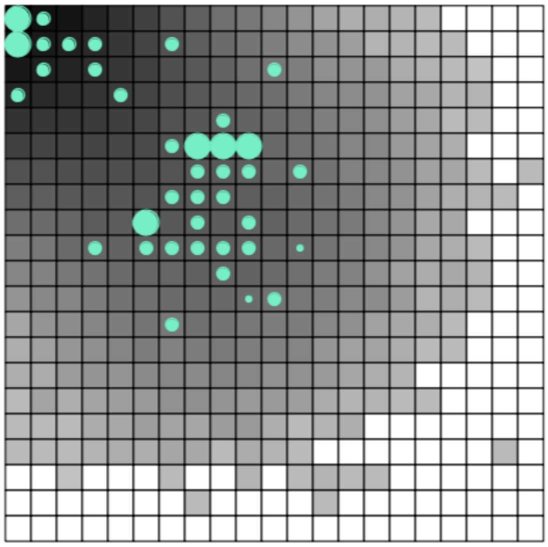

contig_9747000008_Laussat

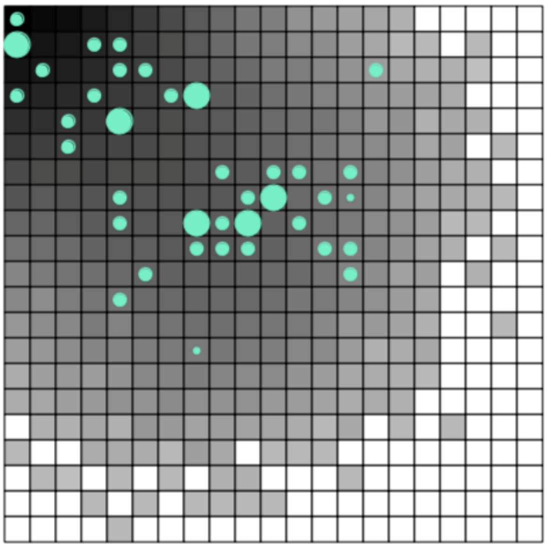

**Comparison of HBM and G2D results**

To check whether the G2D analyses support the HBM results, we compared the distribution of G2D values between HBM outliers and HBM non-outliers, both in the Régina and in the Laussat population pair. The distribution of G2D values is clearly different between the two groups:

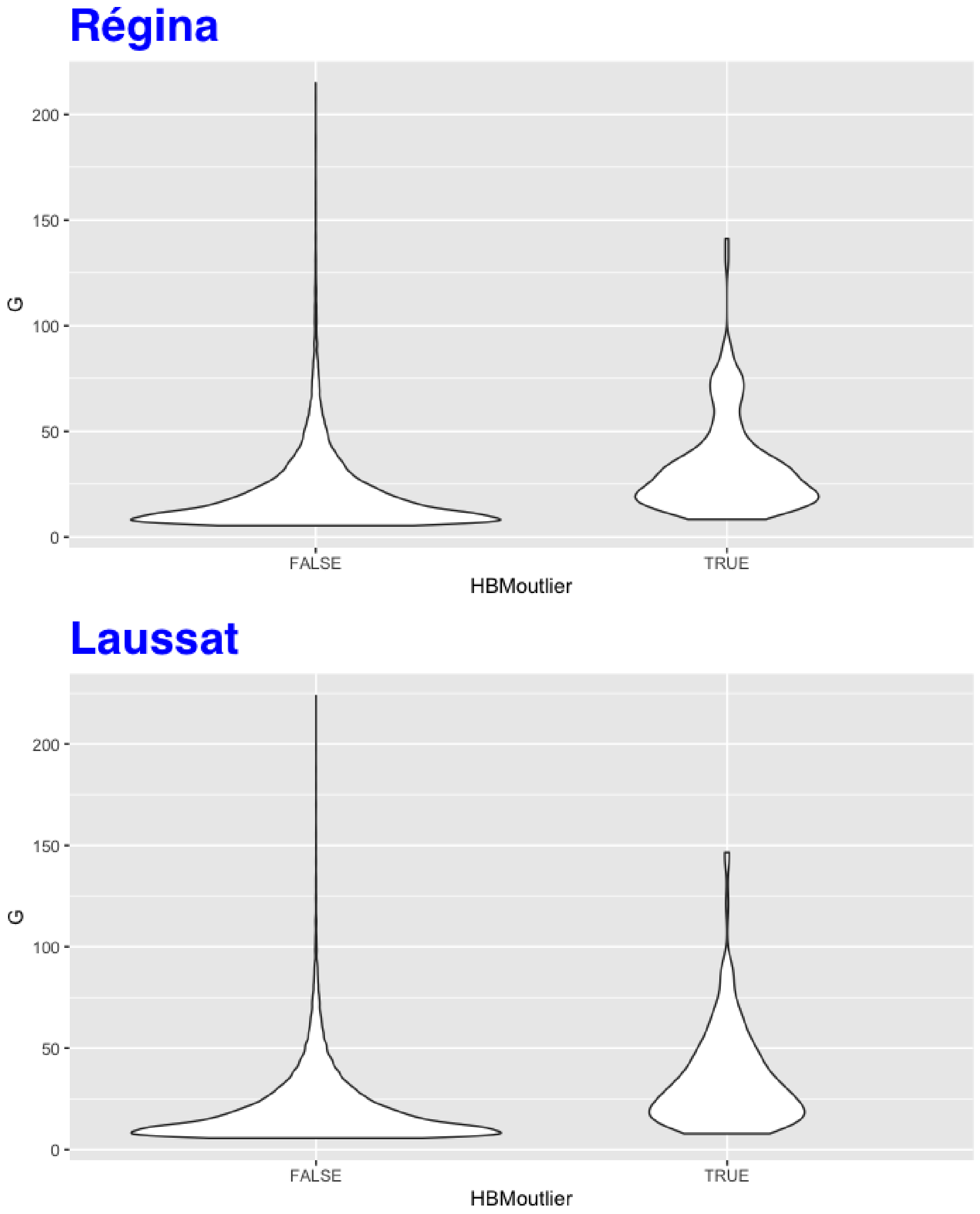


This is confirmed by non-paramatric testing of the differences in G2D values between the outlier and non-outlier groups: Wilcoxon rank sum test with continuity correction for Régina: W = 3106662, p-value < 2.2e-16; Wilcoxon rank sum test with continuity correction for Laussat: W = 3089397, p-value < 2.2e-16.

**Appendix A**

```
G2DcalcMultiPop<-function(sfs=multiPopSFS.list,allfreq=minAllFreq.df,
                          firstPop,secondPop,sampleSize,nPop = 4)
  {
#allows one to compute 2D-SFS for a pair of populations out of a group
#of populations.
#NB this is for PAIRWISE SFS's, not for multi-pop SFS's
#the arguments are:
#allfreq, which defaults to minAllFreq.df
#sfs, which defaults to multiPopSFS.list
#both defaults are inherited from the fromArpToSFS() function
#firstPop and secondPop, the ordinal index of the populations
#sampleSize, the sample per pop
#nPop = number of populations
#EXTRACTING MULTINOMIAL PROBS BY CONTIG
#First, obtaining contingency tables by contig
#writing ad-hoc function
#NB coerce table to sampleSize by sampleSize
#2D-SFS tables are made from allele counts in each pop
#
#levels are coerced to 0:sampleSize (=nb of alleles)
#each table takes a vector form (?) so that the output is a matrix
#with each contig in a column
contigTable<-function(y){table(factor(y[,(firstPop+1)],levels=0:sampleSize),
    factor(y[,(secondPop+1)],levels=0:sampleSize))
  }
#applying by contig (using by())
#INDICES is the contig column in the MinAllFreq.df object:
#it must be written explicitly as an argument (this is normal)
contig2DSFS<-by(minAllFreq.df,FUN=contigTable,INDICES=as.factor(minAllFreq.df[,1]))
#
#implementing multinomial prob calculation for tables:
#the function here takes each element of the matrix-containing list,
#turns them into vectors and computes probabilities
#based on the expectation contained in the global (genome-wide) MAF 2DSFS
#this requires that the genome-wide SFS's name is HARDWIRED
#(I could change this, too)
#
#***IMPORTANT***
#the MAF2DSFS here must be obtained from the SIMULATED DATA
#that have been simulated based on the observed SFS, which
#is NOT USED ANYMORE here.
#first, the right element of multiPopSFS.list must be identified
pointer<-matrix(ncol=2)
for (i in 1:(nPop-1))
{
  for (j in (i+1):nPop)
  {
    pointer<-rbind(pointer,c(i,j))
  }
}
pointer<-pointer[-1,]
pointer<-cbind(pointer,1:nrow(pointer))
sfsIndex<-pointer[pointer[,1]==firstPop & pointer[,2]==secondPop,3]
multinomProbs<-function(X){dmultinom(x=as.vector(X),
                                     prob=as.vector(as.matrix(sfs[[sfsIndex]])))}
#notice that the function is run through a lapply()
#because the starting object is a LIST
ProbsGlobal.list<-lapply(X=contig2DSFS,FUN=multinomProbs)
```

```
#same as above, but computes probs relative to the
#LOCAL expectation
multinomProbsLocal<-function(X){dmultinom(x=as.vector(X),
                                          prob=as.vector(X))}
ProbsLocal.list<-lapply(X=contig2DSFS,FUN=multinomProbsLocal)
#computation of the statistics
Gvalues<<-2*(log(unlist(ProbsLocal.list)) - log(unlist(ProbsGlobal.list)))
#removing uninformative loci (must look at it more closely)
#GvaluesInformative<-Gvalues[which(CROPsProbsDenom.list!=1)]
hist(Gvalues)
# #not used: computation of theoretical P-values
# #P-values: the test must have Ne d.f.
# #(Ne=the number of non-empty cells in the genome-wide table)
# Ne<-length(sfs[sfs!=0])
# #provides the emirical P-value distribution
# #(for more precision, this must be cumulated with several other
# #simulations)
# #not used: vector of P values
# #as.vector(pchisq(q=Gvalues, df=Ne, ncp = 0, lower.tail = TRUE, log.p = FALSE))
Gvalues
#as.vector(Gvalues)
}

#not used: test of G2calc
# firstSetOfGvalues<-G2Dcalc(allfreq=minAllFreq.df,sfs=MAF2DSFS)
```
