## Supplementary material for "Genomic and phenotypic divergence unveil microgeographic adaptation in the Amazonian hyperdominant tree *Eperua falcata* Aubl. (Fabaceae)": File S6

### Reciprocal Transplants methodology

Seeds were sampled around fructifying mother trees within each site and microhabitat according to a grid layout. Seeds were weighted and sown in individual pots containing forest soils in a shade-house daily watered until cotyledons emerged (about one month after seeds sowing). Then, about 800 vigorous seedlings were transplanted into twelve experimental gardens of dimension 10m x 10m located in the undisturbed forests of Laussat and Regina (three gardens per site and microhabitat within each site), with randomization between and within gardens. Seedlings survival, growth (height, diameter, number of growth units, number of leaves and leaflets), leaf properties (leaf area and chlorophyll content apprehended with chlorophyll meter SPAD-502) and predation (in terms of number of leaflets attacked) were measured from September 2011 to September 2014.

Seedlings phenotypic values were partitioned into genetic and environmental effects at both the regional and the microgeographic scale according to a classical linear model of quantitative genetics, by imposing the 'zero-sum' constraint for each factor. Four different models were compared (with and without interactions, and with and without the garden as a random effect) by retaining the model with the lowest Akaike information criterion (AIC). Seed mass was incorporated as a quantitative cofactor to each model.

<u>Model 1 (full model):</u> Pi = μ + SMi + RPi + RTi + I(RPi,RTi) + MPi + MTi + I(MPi,MTi) + (1|Gi) + εi

where μ is the global mean, SM is the fresh seed mass, RP is the regional provenance or site of provenance, RT is the regional site of transplantation, MP is the microgeographic environment of provenance, MT is the microgeographic environment of transplantation, G the garden of transplantation (random effect) and ε the residuals.

<u>Model 2 (without interactions):</u> Pi = μ + SMi + RPi + RTi + MPi + MTi + (1|Gi) + εi

<u>Model 3 (without random effect):</u> Pi = μ + SMi + RPi + RTi + I(RPi,RTi) + MPi + MTi + I(MPi,MTi) + εi

<u>Model 4 (without interactions nor random effect):</u> Pi = μ + SMi + RPi + RTi + MPi + MTi + + εi

Models 1 and 3 were fitted with the functions 'glmer' and 'lmer' for discrete (survival and discrete counts) and continuous (height, diameter) traits respectively, while models 2 and 4 were fitted with the function 'glm'. A binomial distribution was used for binary traits (survival), a Poisson distribution was used for discrete counts, and a Gaussian distribution was used for continuous traits.
