## Supplementary material for "Genomic and phenotypic divergence unveil microgeographic adaptation in the Amazonian hyperdominant tree *Eperua falcata* Aubl. (Fabaceae)": File S7

### Bioinformatics results

The crude reference genome produced here, although incomplete and fragmented, constitutes a new, valuable pan-genomic resource containing key information about population-level genetic diversity (i.e. tens of thousands of loci scattered throughout the genome, a first for a non-model tree species of Amazonia). The four population pools were assembled *de novo* into a high-coverage pan-genomic reference: *de novo* assembly led to 325,249 contigs in 272,865 scaffolds for a total assembly length of about 250 Mb (252,364,908 bases assembled *de novo* of which 249,308,901 were successfully mapped *a posteriori* by the different libraries, **Table S1**) and a mapping coverage of 57.81X on average, **Fig. S3**. The structural annotation predicted 32,789 genes and 117,278 exons, and detected 124,287 repeated elements. A total of 32,075 predicted genes were successfully annotated using BLAST, and only 732 predicted genes did not return any BLAST hits. Additionally, gene-ontology terms (GO terms) were retrieved for 18,577 predicted genes, while the remaining 14,212 genes were not associated to any GO term.
