## Supplementary material for "Genomic and phenotypic divergence unveil microgeographic adaptation in the Amazonian hyperdominant tree *Eperua falcata* Aubl. (Fabaceae)": File S8

| | geneID | sacc | descr | GO_CHILDS_AND_PARENTS_NAMES | scaffold | start | end | N_outliers_loc | Enrichment te | Subcellular loc | Link | comment |
|---|---|---|---|---|---|---|---|---|---|---|---|---|
| 546 | predgene_003755 | OMO49514.1 | hypothetical protein CCACVL1_30963 [Corchorus capsularis] | NA | scaffold-1039 | 166 | 6421 | 1 | no | | | NA |
| 555 | predgene_000745 | XP_020224053.1 | probable cellulose synthase A catalytic subunit 3 [UDP-forming] isoform X2 [Cajanus cajan] | membrane;cellulose synthase (UDP-forming) activity;cellulose biosynthe | scaffold-1040 | 955 | 9058 | 1 | yes | | http://www.uniprot.org/uniparc/UPI00098D75F7 | stress, signalization and transport, regulation |
| 574 | predgene_018831 | KRG93973.1 | hypothetical protein GLYMA_19G053500 [Glycine max] | glycerol-3-phosphate dehydrogenase [NAD+] activity;glycerol-3-phospha | scaffold-10411 | 61 | 1648 | 4 | yes | | http://www.uniprot.org/uniprot/I1N6W7 | primary processes, stress, armoatic compounds |
| 753 | predgene_027371 | AFK45765.1 | unknown [Lotus japonicus] | metabolic process;methyltransferase activity;biological_process;transfer | scaffold-105544 | 2 | 968 | 1 | yes | | | NA |
| 757 | predgene_021333 | XP_015577436.1 | PREDICTED: phenylalanine N-monooxygenase-like [Ricinus communis] | iron ion binding;oxidoreductase activity, acting on paired donors, with in | scaffold-10558 | 361 | 1379 | 1 | yes | | http://www.uniprot.org/uniparc/UPI000772731A | primary processes, stress, armoatic compounds |
| 1122 | predgene_006069 | KJB25052.1 | hypothetical protein B456_004G179200 [Gossypium raimondii] | NA | scaffold-10829 | 2935 | 5236 | 1 | no | | | NA |
| 1236 | predgene_006075 | XP_020239235.1 | palmitoyl-acyl carrier protein thioesterase, chloroplastic-like isoform X1 [Cajanus cajan] | fatty acid biosynthetic process;thiolester hydrolase activity;fatty acid me | scaffold-10923 | 1741 | 5276 | 1 | yes | | http://www.uniprot.org/uniparc/UPI00098D8922 | lipid metabolism |
| 1722 | predgene_006329 | XP_020257616.1 | diacylglycerol kinase 1-like isoform X3 [Asparagus officinalis] | diacylglycerol kinase activity;protein kinase C-activating G-protein couple | scaffold-11331 | 610 | 5446 | 1 | yes | | http://www.uniprot.org/uniparc/UPI00098DFD96 | primary processes, stress, signalization and transport, regulation |
| 1741 | predgene_000062 | XP_006592370.1 | PREDICTED: dirigent protein 2-like [Glycine max] | NA | scaffold-11345 | 16334 | 17142 | 1 | no | | | NA |
| 1742 | predgene_000058 | XP_010657210.1 | PREDICTED: DNA repair protein RAD16 [Vitis vinifera] | ATP binding;adenyl ribonucleotide binding;purine ribonucleoside triphos | scaffold-11345 | 401 | 4947 | 1 | yes | | http://www.uniprot.org/uniprot/F6HIC3 | primary processes, armoatic compounds |
| 1743 | predgene_000061 | XP_003605958.1 | B-cell receptor-associated-like protein [Medicago truncatula] | endoplasmic reticulum;intracellular protein transport;integral componer | scaffold-11345 | 14559 | 14864 | 1 | yes | endoplasmic r | http://www.uniprot.org/uniprot/G7JEQ0 | primary processes, signalization and transport, regulation |
| 1744 | predgene_000060 | XP_004506274.1 | PREDICTED: B-cell receptor-associated protein 31 [Cicer arietinum] | endoplasmic reticulum;intracellular protein transport;integral componer | scaffold-11345 | 11657 | 12194 | 1 | yes | endoplasmic r | http://www.uniprot.org/uniprot/A0A1S2YL36 | primary processes, signalization and transport, regulation |
| 1745 | predgene_000059 | XP_003541950.1 | PREDICTED: DNA repair protein RAD16-like isoform X3 [Glycine max] | metal ion binding;protein binding;zinc ion binding;cation binding;binding | scaffold-11345 | 6948 | 9909 | 1 | no | | http://www.uniprot.org/uniprot/K7M2X2 | stress, regulation |
| 1746 | predgene_000063 | XP_019451198.1 | PREDICTED: mitogen-activated protein kinase kinase kinase A-like [Lupinus angustifolius] | protein kinase activity;protein phosphorylation;ATP binding;kinase activi | scaffold-11345 | 19685 | 21821 | 1 | yes | | http://www.uniprot.org/uniprot/A0A1J7HW07 | primary processes, armoatic compounds, regulation |
| 2753 | predgene_010285 | XP_016542622.1 | PREDICTED: serine/threonine-protein kinase AFC1 isoform X1 [Capsicum annuum] | protein kinase activity;ATP binding;protein phosphorylation;kinase activi | scaffold-12168 | 825 | 3807 | 1 | yes | | http://www.uniprot.org/uniprot/A0A1U8E5V2 | primary processes, armoatic compounds, regulation |
| 2769 | predgene_003759 | KJB65030.1 | hypothetical protein B456_010G077100 [Gossypium raimondii] | NA | scaffold-1218 | 539 | 5652 | 1 | no | | | NA |
| 3051 | predgene_015938 | XP_003621392.2 | hydrolyzing O-glycosyl compounds hydrolase [Medicago truncatula] | hydrolase activity, hydrolyzing O-glycosyl compounds;carbohydrate met | scaffold-124136 | 740 | 2709 | 1 | yes | | | NA |
| 3638 | predgene_007379 | AKF43194.1 | metallopeptidase M24 family protein [Francoa sonchifolia] | NA | scaffold-12903 | 577 | 4970 | 2 | no | | | NA |
| 3665 | predgene_005012 | OIW07762.1 | hypothetical protein TanjilG_12888 [Lupinus angustifolius] | protein binding;binding;molecular_function | scaffold-12921 | 104 | 1860 | 1 | no | | http://www.uniprot.org/uniprot/A0A1J7HMW3 | regulation |
| 3666 | predgene_005013 | XP_007139804.1 | hypothetical protein PHAVU_008G060200g [Phaseolus vulgaris] | NA | scaffold-12921 | 2280 | 4178 | 1 | no | | | NA |
| 4494 | predgene_008954 | XP_015955508.1 | probable E3 ubiquitin-protein ligase RHB1A [Arachis duranensis] | protein binding;zinc ion binding;binding;transition metal ion binding;mol | scaffold-13610 | 1209 | 2679 | 1 | no | | http://www.uniprot.org/uniparc/UPI000786E800 | stress, regulation |
| 5046 | predgene_008697 | XP_012575122.1 | PREDICTED: protein-lysine methyltransferase METTL21B isoform X2 [Cicer arietinum] | NA | scaffold-140674 | 409 | 3683 | 1 | no | | | NA |
| 5088 | predgene_004360 | XP_012567275.1 | PREDICTED: U6 snRNA phosphodiesterase isoform X4 [Cicer arietinum] | nuclease activity;U6 snRNA 3'-end processing;hydrolase activity, acting o | scaffold-14107 | 2690 | 4436 | 1 | yes | nucleus | http://www.uniprot.org/uniprot/A0A1S3DV08 | primary processes, armoatic compounds, expression |
| 5089 | predgene_004361 | KHN30056.1 | hypothetical protein glysoja_010434 [Glycine soja] | NA | scaffold-14107 | 5949 | 6461 | 1 | no | | | NA |
| 5090 | predgene_004359 | XP_020224625.1 | heterogeneous nuclear ribonucleoprotein 1 isoform X2 [Cajanus cajan] | nucleotide binding;nucleic acid binding;small molecule binding;nucleosid | scaffold-14107 | 244 | 1323 | 1 | yes | | http://www.uniprot.org/uniparc/UPI00098DAA44 | primary processes, armoatic compounds, expression |
| 5286 | predgene_000254 | XP_014503952.1 | acidic mammalian chitinase [Vigna radiata var. radiata] | hydrolase activity, hydrolyzing O-glycosyl compounds;carbohydrate met | scaffold-14311 | 748 | 2054 | 1 | yes | | http://www.uniprot.org/uniprot/A0A1S3UD38 | primary processes, stress |
| 5287 | predgene_000255 | XP_020213761.1 | CHD3-type chromatin-remodeling factor PICKLE isoform X1 [Cajanus cajan] | NA | scaffold-14311 | 4232 | 5488 | 1 | no | | | NA |
| 5288 | predgene_000256 | XP_014500994.1 | CHD3-type chromatin-remodeling factor PICKLE isoform X3 [Vigna radiata var. radiata] | DNA binding;ATP binding;nucleus;regulation of transcription, DNA-templ | scaffold-14311 | 6633 | 13203 | 1 | yes | nucleus | http://www.uniprot.org/uniprot/A0A1S3U4S4 | primary processes, stress, signalization and transport, armoatic compounds, expression, regulation |
| 5976 | predgene_004515 | OMO70780.1 | Tetratricopeptide-like helical [Corchorus olitorius] | protein binding;binding;molecular_function | scaffold-14924 | 2433 | 5362 | 1 | no | | http://www.uniprot.org/uniprot/A0A1R3HKD7 | regulation |
| 6002 | predgene_003732 | XP_020226459.1 | histone-lysine N-methyltransferase EZA1-like isoform X2 [Cajanus cajan] | protein binding;histone-lysine N-methyltransferase activity;seed develop | scaffold-14952 | 618 | 6616 | 1 | yes | | http://www.uniprot.org/uniparc/UPI00098DADE5 | primary processes, development, reproduction, expression, regulation |
| 7050 | predgene_009237 | XP_019184053.1 | PREDICTED: nuclear transport factor 2 [Ipomoea nil] | intracellular;transport;cell part;establishment of localization;cellular_cor | scaffold-15978 | 3715 | 4334 | 1 | no | | http://www.uniprot.org/uniparc/UPI00090177E9 | signalization and transport |
| 7085 | predgene_005799 | XP_016463998.1 | PREDICTED: ubiquitin carboxyl-terminal hydrolase 4-like isoform X1 [Nicotiana tabacum] | ubiquitin-dependent protein catabolic process;thiol-dependent ubiquitin | scaffold-16011 | 208 | 5228 | 1 | yes | | http://www.uniprot.org/uniprot/A0A1S3ZHT9 | primary processes, regulation |
| 7205 | predgene_006243 | OWM78043.1 | hypothetical protein CDL15_Pgr018612 [Punica granatum] | nucleotide binding;nucleic acid binding;small molecule binding;nucleosid | scaffold-16122 | 947 | 5202 | 1 | yes | | http://www.uniprot.org/uniprot/A0A218WZF5 | primary processes, armoatic compounds, expression |
| 7317 | predgene_012747 | KHN45096.1 | DUF246 domain-containing protein [Glycine soja] | NA | scaffold-16235 | 817 | 3259 | 1 | no | | | NA |
| 7492 | predgene_019500 | XP_006581972.1 | PREDICTED: GDSL esterase/lipase At1g74460-like [Glycine max] | hydrolase activity, acting on ester bonds;hydrolase activity;catalytic activ | scaffold-16406 | 3 | 1247 | 1 | yes | | | NA |
| 7581 | predgene_016683 | XP_007153735.1 | hypothetical protein PHAVU_003G060700g [Phaseolus vulgaris] | protein dimerization activity;protein binding;binding;molecular_function | scaffold-16494 | 2 | 1622 | 1 | no | | http://www.uniprot.org/uniprot/V7C6I7 | regulation |
| 7657 | predgene_014420 | XP_020221813.1 | uncharacterized protein LOC109804406 [Cajanus cajan] | NA | scaffold-16551 | 1 | 983 | 1 | no | | | NA |
| 7868 | predgene_024205 | XP_015882433.1 | PREDICTED: adenosylhomocysteinase 1-like [Ziziphus jujuba] | adenosylhomocysteinase activity;S-adenosylhomocysteine catabolic pro | scaffold-167762 | 205 | 957 | 1 | yes | | http://www.uniprot.org/uniparc/UPI00077E9004 | primary processes, armoatic compounds |
| 8448 | predgene_000677 | OAP11746.1 | hypothetical protein AXX17_AT2G33830 [Arabidopsis thaliana] | protein kinase activity;protein phosphorylation;ATP binding;protein bind | scaffold-17377 | 1659 | 10579 | 1 | yes | membrane | http://www.uniprot.org/uniprot/A0A178W0X1 | primary processes, armoatic compounds, regulation |
| 8859 | predgene_011829 | KMZ58531.1 | vascular plant one zinc finger protein [Zostera marina] | NA | scaffold-17796 | 852 | 3749 | 1 | no | | | NA |
| 9241 | predgene_001856 | EFH61567.1 | hypothetical protein ARALYDRAFT_342068 [Arabidopsis lyrata subsp. lyrata] | NA | scaffold-18218 | 183 | 8064 | 1 | no | | | NA |
| 9646 | predgene_005648 | XP_021685116.1 | uncharacterized protein LOC110668256 isoform X2 [Hevea brasiliensis] | folic acid binding;metabolic process;transferase activity;vitamin binding; | scaffold-18667 | 171 | 964 | 1 | yes | | http://www.uniprot.org/uniparc/UPI000B775932 | armoatic compounds |
| 9647 | predgene_005649 | XP_022634329.1 | beta-glucuronosyltransferase GlcAT14B isoform X1 [Vigna radiata var. radiata] | acetylglucosaminyltransferase activity;membrane;UDP-glycosyltransfera | scaffold-18667 | 1242 | 4148 | 1 | yes | | http://www.uniprot.org/uniparc/UPI000BE42773 | signalization and transport |
| 9718 | predgene_010056 | XP_012571111.1 | PREDICTED: LOW QUALITY PROTEIN: protein SUPPRESSOR OF npr1-1, CONSTITUTIVE 1-like [Cicer arietinum] | NA | scaffold-18765 | 3594 | 3797 | 1 | no | | | NA |
| 9901 | predgene_001115 | XP_020089007.1 | uncharacterized protein LOC109710684 [Ananas comosus] | NA | scaffold-18962 | 993 | 1268 | 1 | no | | | NA |
| 10284 | predgene_012247 | KVI01320.1 | hypothetical protein Ccrd_020418 [Cynara cardunculus var. scolymus] | oxoglutarate dehydrogenase (succinyl-transferring) activity;tricarboxylic | scaffold-194 | 523 | 3304 | 1 | yes | | http://www.uniprot.org/uniprot/A0A118K0S7 | primary processes, stress, armoatic compounds |
| 11165 | predgene_017213 | XP_014628904.1 | PREDICTED: acyl-coenzyme A thioesterase 8 isoform X2 [Glycine max] | acyl-CoA metabolic process;acyl-CoA hydrolase activity;coenzyme metab | scaffold-20365 | 233 | 2701 | 1 | yes | | | NA |
| 11723 | predgene_006300 | XP_015954463.1 | DNA repair protein recA homolog 3, mitochondrial isoform X2 [Arachis duranensis] | single-stranded DNA binding;ATP binding;DNA repair;DNA binding;DNA r | scaffold-21020 | 507 | 5556 | 1 | yes | | http://www.uniprot.org/uniparc/UPI0007896541 | primary processes, stress, signalization and transport, armoatic compounds, expression |
| 12218 | predgene_002714 | ACN85301.1 | putative RNA polymerase A(I) large subunit [Oryza coarctata] | DNA-directed 5'-3' RNA polymerase activity;nucleus;transcription, DNA-t | scaffold-21679 | 609 | 7566 | 1 | yes | nucleus | http://www.uniprot.org/uniprot/C0JAC0 | primary processes, stress, signalization and transport, armoatic compounds, expression |
| 12248 | predgene_003398 | OEL36438.1 | DEAD-box ATP-dependent RNA helicase 8 [Dichanthelium oligosanthes] | nucleic acid binding;ATP binding;organic cyclic compound binding;heterc | scaffold-21735 | 1286 | 4390 | 1 | yes | membrane | http://www.uniprot.org/uniprot/A0A1E5WGF2 | primary processes, armoatic compounds, expression |
| 12249 | predgene_003399 | XP_017188591.1 | PREDICTED: DEAD-box ATP-dependent RNA helicase 8-like [Malus domestica] | NA | scaffold-21735 | 5787 | 6482 | 1 | no | | | NA |
| 12294 | predgene_010229 | XP_010103434.1 | putative serine incorporator [Morus notabilis] | membrane;cellular_component | scaffold-2180 | 264 | 3920 | 1 | no | transmembrai | http://www.uniprot.org/uniprot/W9RU03 | signalization and transport |
| 12741 | predgene_002577 | XP_010682358.1 | PREDICTED: oxysterol-binding protein-related protein 1D [Beta vulgaris subsp. vulgaris] | NA | scaffold-22355 | 1052 | 6904 | 1 | no | | | NA |
| 12742 | predgene_002578 | GAU46157.1 | hypothetical protein TSUD_301170 [Trifolium subterraneum] | NA | scaffold-22355 | 7260 | 7442 | 1 | no | | | NA |
| 12927 | predgene_001066 | KHN43438.1 | Long chain acyl-CoA synthetase 1 [Glycine soja] | catalytic activity;metabolic process;molecular_function;biological_proce | scaffold-22573 | 3931 | 8857 | 1 | yes | | | NA |
| 13253 | predgene_020657 | XP_020973327.1 | geranylgeranyl transferase type-2 subunit beta 1 isoform X3 [Arachis ipaensis] | Rab geranylgeranyltransferase activity;protein geranylgeranylation;catal | scaffold-229908 | 261 | 1487 | 1 | yes | | http://www.uniprot.org/uniparc/UPI000A2B13B9 | primary processes, regulation |
| 13602 | predgene_006705 | XP_020236842.1 | serine/arginine repetitive matrix protein 1 [Cajanus cajan] | NA | scaffold-234430 | 1916 | 5300 | 1 | no | | | NA |
| 13949 | predgene_028120 | OMO92626.1 | Cytochrome P450 [Corchorus olitorius] | NA | scaffold-239179 | 87 | 353 | 1 | no | | | NA |
| 14066 | predgene_011486 | XP_009758015.1 | PREDICTED: copper-transporting ATPase RAN1-like, partial [Nicotiana sylvestris] | integral component of membrane;metal ion transport;metal ion binding | scaffold-24077 | 3 | 3746 | 1 | yes | transmembrai | http://www.uniprot.org/uniprot/A0A1U7V8A1 | primary processes, stress, signalization and transport, armoatic compounds |
| 14922 | predgene_007023 | XP_019413900.1 | PREDICTED: delta-1-pyrroline-5-carboxylate synthase-like isoform X2 [Lupinus angustifolius] | metabolic process;oxidoreductase activity;oxidation-reduction process;o | scaffold-25260 | 283 | 3184 | 1 | yes | cytoplasm | http://www.uniprot.org/uniprot/A0A1J7GDX7 | stress |
| 14923 | predgene_007024 | XP_014633475.1 | PREDICTED: delta-1-pyrroline-5-carboxylate synthase-like isoform X4 [Glycine max] | metabolic process;oxidoreductase activity;oxidation-reduction process;g | scaffold-25260 | 3348 | 4879 | 1 | yes | | http://www.uniprot.org/uniprot/K7L1J5 | primary processes, stress, armoatic compounds |
| 15145 | predgene_008667 | NA | NA | NA | scaffold-2556 | 3041 | 3103 | 1 | no | | | NA |
| 15240 | predgene_002974 | XP_015934152.1 | lysM domain receptor-like kinase 4 [Arachis duranensis] | protein kinase activity;ATP binding;protein phosphorylation;kinase activi | scaffold-25703 | 1517 | 3412 | 1 | yes | | http://www.uniprot.org/uniparc/UPI0007899EF5 | primary processes, armoatic compounds, regulation |
| 15241 | predgene_002975 | GAU15346.1 | hypothetical protein TSUD_04150 [Trifolium subterraneum] | zinc ion binding;transition metal ion binding;metal ion binding;cation bin | scaffold-25703 | 3599 | 7086 | 1 | no | NA | NA | stress |
| 16360 | predgene_012189 | XP_019440191.1 | PREDICTED: uncharacterized protein LOC109345566 [Lupinus angustifolius] | NA | scaffold-2717 | 114 | 3189 | 1 | no | | | NA |
| 16399 | predgene_014347 | XP_016447680.1 | PREDICTED: metal tolerance protein 4-like [Nicotiana tabacum] | cation transport;cation transmembrane transporter activity;integral com | scaffold-27225 | 575 | 1885 | 2 | no | transmembrai | http://www.uniprot.org/uniprot/A0A1S3Y6E8 | signalization and transport |
| 16473 | predgene_013506 | XP_004488935.1 | PREDICTED: uncharacterized membrane protein At3g27390-like [Cicer arietinum] | NA | scaffold-272753 | 83 | 2341 | 1 | no | | | NA |
| 17104 | predgene_021059 | XP_003589223.1 | hypothetical protein MTR_1g019800 [Medicago truncatula] | NA | scaffold-28290 | 648 | 1612 | 1 | no | | | NA |
| 17876 | predgene_000563 | XP_010246915.1 | PREDICTED: centriolin isoform X2 [Nelumbo nucifera] | NA | scaffold-30110 | 6776 | 10781 | 1 | no | | | NA |
| 18008 | predgene_019003 | XP_018812405.1 | PREDICTED: laccase-7-like [Juglans regia] | copper ion binding;oxidoreductase activity;oxidation-reduction process;t | scaffold-30393 | 553 | 1302 | 1 | yes | | http://www.uniprot.org/uniparc/UPI0008DD5096 | stress |
| 18305 | predgene_008605 | XP_022151653.1 | uncharacterized protein LOC111019565 [Momordica charantia] | NA | scaffold-31051 | 410 | 769 | 1 | no | | | NA |
| 18514 | predgene_011170 | KHN36487.1 | mRNA-decapping enzyme-like protein [Glycine soja] | deadenylation-dependent decapping of nuclear-transcribed mRNA;enzyr | scaffold-31599 | 1838 | 4034 | 1 | yes | | http://www.uniprot.org/uniparc/UPI00054A00F9 | primary processes, armoatic compounds, expression, regulation |
| 18742 | predgene_015447 | XP_008341863.1 | PREDICTED: uncharacterized protein LOC103404704 [Malus domestica] | NA | scaffold-32177 | 2268 | 2408 | 1 | no | | | NA |
| 19159 | predgene_009993 | KJB83145.1 | hypothetical protein B456_013G231400 [Gossypium raimondii] | membrane;cellular_component | scaffold-32933 | 65 | 3950 | 1 | no | transmembrai | http://www.uniprot.org/uniprot/A0A0D2W8Q4 | signalization and transport |
| 19497 | predgene_011112 | XP_014502347.1 | plant intracellular Ras-group-related LRR protein 9 [Vigna radiata var. radiata] | protein binding;binding;molecular_function | scaffold-336 | 3 | 2993 | 1 | no | | http://www.uniprot.org/uniprot/A0A1S3U8J9 | regulation |
| 19498 | predgene_011113 | KYP69632.1 | Protein lap1, partial [Cajanus cajan] | protein binding;binding;molecular_function | scaffold-336 | 3047 | 3556 | 1 | no | | http://www.uniprot.org/uniprot/A0A151TRE0 | regulation |
| 20100 | predgene_010110 | XP_006444243.1 | hypothetical protein CICLE_v10023999mg [Citrus clementina] | electron carrier activity;protein disulfide oxidoreductase activity;cell red | scaffold-350 | 2728 | 3036 | 1 | yes | | http://www.uniprot.org/uniprot/V4TAX8 | primary processes, stress, regulation |
| 20565 | predgene_017715 | XP_003626360.2 | ripening-regulated protein [Medicago truncatula] | NA | scaffold-36134 | 232 | 1761 | 1 | no | | | NA |
| 20959 | predgene_018494 | XP_012089573.1 | THO complex subunit 4B [Jatropha curcas] | nucleic acid binding;nucleotide binding;organic cyclic compound binding; | scaffold-3721 | 206 | 1243 | 1 | yes | | http://www.uniprot.org/uniprot/A0A067L956 | primary processes, armoatic compounds, expression |
| 21147 | predgene_019656 | OMO87526.1 | Clathrin adaptor, mu subunit [Corchorus capsularis] | intracellular protein transport;vesicle-mediated transport;clathrin adapt | scaffold-37785 | 154 | 2278 | 1 | yes | | http://www.uniprot.org/uniprot/A0A1R3IY77 | primary processes, signalization and transport, regulation |
| 21290 | predgene_012205 | EOY23337.1 | DNA primases isoform 3 [Theobroma cacao] | NA | scaffold-38266 | 8 | 3612 | 1 | no | | | NA |
| 22326 | predgene_005668 | XP_009376238.1 | PREDICTED: probable ubiquitin-like-specific protease 2B isoform X5 [Pyrus x bretschneideri] | proteolysis;cysteine-type peptidase activity;protein metabolic process;p | scaffold-41601 | 1009 | 5464 | 1 | yes | | http://www.uniprot.org/uniparc/UPI000510AA8D | primary processes, regulation |
| 23100 | predgene_011096 | ONM39171.1 | 65-kDa microtubule-associated protein 3 [Zea mays] | microtubule cytoskeleton organization;cytokinesis;microtubule binding;c | scaffold-44553 | 1 | 2763 | 1 | yes | | http://www.uniprot.org/uniprot/A0A1D6NFB7 | regulation |
| 23112 | predgene_016728 | XP_010089088.1 | Mitogen-activated protein kinase kinase kinase 2 [Morus notabilis] | NA | scaffold-4462 | 74 | 565 | 1 | no | | | NA |
| 23113 | predgene_016729 | XP_020216595.1 | short-chain dehydrogenase TIC 32, chloroplastic isoform X2 [Cajanus cajan] | NA | scaffold-4462 | 1808 | 2788 | 1 | no | | | NA |
| 23890 | predgene_002382 | XP_012485266.1 | PREDICTED: vacuolar protein sorting-associated protein 26A-like [Gossypium raimondii] | NA | scaffold-4778 | 662 | 6244 | 1 | no | | | NA |
| 24554 | predgene_020677 | GAU30267.1 | hypothetical protein TSUD_384880 [Trifolium subterraneum] | NA | scaffold-50627 | 358 | 519 | 1 | no | | | NA |
| 24632 | predgene_009215 | XP_006573922.1 | PREDICTED: 1-aminocyclopropane-1-carboxylate synthase 7-like [Glycine max] | biosynthetic process;pyridoxal phosphate binding;catalytic activity;meta | scaffold-50996 | 1 | 1051 | 1 | yes | | http://www.uniprot.org/uniprot/I1J4D1 | primary processes, armoatic compounds |
| 25473 | predgene_009529 | OMO61229.1 | Leucine-rich repeat, ribonuclease inhibitor subtype [Corchorus capsularis] | protein binding;binding;molecular_function | scaffold-55 | 577 | 2190 | 1 | no | | http://www.uniprot.org/uniprot/A0A1R3GT44 | regulation |
| 25634 | predgene_013852 | APR64411.1 | hypothetical protein [Populus tomentosa] | NA | scaffold-55898 | 190 | 681 | 1 | no | | | NA |
| 25635 | predgene_013853 | CDO97347.1 | unnamed protein product [Coffea canephora] | protein binding;zinc ion binding;binding;transition metal ion binding;mol | scaffold-55898 | 2215 | 3378 | 1 | no | | http://www.uniprot.org/uniprot/A0A068TMV6 | stress, regulation |
| 26249 | predgene_000689 | XP_020218813.1 | phosphatidylinositol/phosphatidylcholine transfer protein SFH9 isoform X3 [Cajanus cajan] | NA | scaffold-5909 | 3725 | 10323 | 1 | no | | | NA |
| 26250 | predgene_000688 | KRG99755.1 | hypothetical protein GLYMA_18G169200 [Glycine max] | NA | scaffold-5909 | 148 | 479 | 1 | no | | | NA |
| 26805 | predgene_012047 | KVH91672.1 | Adaptor protein complex AP-4, epsilon subunit [Cynara cardunculus var. scolymus] | binding;intracellular protein transport;vesicle-mediated transport;memb | scaffold-61904 | 56 | 3689 | 1 | yes | | http://www.uniprot.org/uniprot/A0A118JUK7 | primary processes, signalization and transport, regulation |
| 27266 | predgene_016594 | XP_020997687.1 | ankyrin repeat domain-containing protein, chloroplastic [Arachis duranensis] | protein binding;binding;molecular_function | scaffold-64285 | 106 | 2695 | 1 | no | | http://www.uniprot.org/uniparc/UPI000A2C2546 | regulation |
| 27647 | predgene_005688 | XP_019072172.1 | PREDICTED: uncharacterized protein LOC100253998 isoform X1 [Vitis vinifera] | NA | scaffold-6633 | 89 | 3415 | 2 | no | | | NA |
| 27763 | predgene_009369 | XP_006859027.1 | protein SPA1-RELATED 4 isoform X1 [Amborella trichopoda] | protein kinase activity;ATP binding;protein phosphorylation;protein bind | scaffold-6692 | 1896 | 4485 | 1 | yes | | http://www.uniprot.org/uniprot/U5DD77 | primary processes, armoatic compounds, regulation |
| 27775 | predgene_003312 | ONM27008.1 | T-complex protein 1 subunit theta [Zea mays] | ATP binding;protein folding;unfolded protein binding;adenyl ribonucleot | scaffold-670 | 144 | 6179 | 1 | yes | cytoplasm | http://www.uniprot.org/uniprot/A0A1D6F6Y7 | primary processes, armoatic compounds, regulation |
| 27985 | predgene_003921 | XP_020236346.1 | plant cysteine oxidase 4-like [Cajanus cajan] | oxidoreductase activity, acting on single donors with incorporation of mc | scaffold-6830 | 1133 | 3198 | 1 | yes | | http://www.uniprot.org/uniparc/UPI00098DC841 | primary processes, stress |
| 29525 | predgene_022002 | XP_021829682.1 | probable calcium-binding protein CML48 [Prunus avium] | calcium ion binding;metal ion binding;cation binding;ion binding;binding; | scaffold-77560 | 3 | 1888 | 1 | no | | http://www.uniprot.org/uniparc/UPI000B7B16AC | stress |
| 30984 | predgene_007301 | XP_012835746.1 | PREDICTED: 5'-3' exoribonuclease 3 isoform X1 [Erythranthe guttata] | nucleic acid binding;exonuclease activity;5'-3' exoribonuclease activity;n | scaffold-8718 | 171 | 5142 | 1 | yes | | http://www.uniprot.org/uniparc/UPI00064D88B9 | primary processes, signalization and transport, armoatic compounds, expression |
| 32313 | predgene_009374 | XP_003592607.1 | DUF789 family protein [Medicago truncatula] | NA | scaffold-9670 | 185 | 3922 | 1 | no | | | NA |
| 32586 | predgene_003725 | XP_020204609.1 | probable serine/threonine-protein kinase PBL7 isoform X2 [Cajanus cajan] | protein kinase activity;ATP binding;protein phosphorylation;kinase activi | scaffold-9862 | 5929 | 6927 | 1 | yes | | http://www.uniprot.org/uniparc/UPI00098D8B10 | primary processes, armoatic compounds, regulation |
| 32591 | predgene_006057 | GAU47782.1 | hypothetical protein TSUD_193190, partial [Trifolium subterraneum] | NA | scaffold-9866 | 771 | 1123 | 1 | no | | | NA |
| 32592 | predgene_006059 | XP_015883548.1 | PREDICTED: uncharacterized protein LOC107419307 [Ziziphus jujuba] | NA | scaffold-9866 | 3281 | 5269 | 1 | no | | | NA |
| 32593 | predgene_006058 | KJB50678.1 | hypothetical protein B456_008G182300 [Gossypium raimondii] | NA | scaffold-9866 | 1477 | 2070 | 1 | no | | | NA |
